# A coarse-grained bacterial cell model for resource-aware analysis and design of synthetic gene circuits

**DOI:** 10.1101/2023.04.08.536106

**Authors:** Kirill Sechkar, Giansimone Perrino, Guy-Bart Stan

## Abstract

Synthetic genes compete among themselves and with the host cell’s genes for expression machinery, exhibiting resource couplings that affect the dynamics of cellular processes. The modeling of such couplings can be facilitated by simplifying the kinetics of resource-substrate binding. Model-guided design allows to counter unwanted indirect interactions by using biomolecular controllers or tuning the biocircuit’s parameters. However, resource-aware biocircuit design in bacteria is complicated by the interdependence of resource availability and cell growth rate, which significantly affects biocircuit performance. This phenomenon can be captured by coarse-grained models of the whole bacterial cell. The level of detail in these models must balance accurate representation of metabolic regulation against model simplicity and interpretability.

We propose a coarse-grained *E. coli* cell model that combines the ease of simplified resource coupling analysis with the appreciation of bacterial growth regulation mechanisms. Reliably capturing known growth phenomena, it enables numerical prototyping of biocircuits and derivation of analytical relations which can guide the design process. By reproducing several distinct empirical laws observed in prior studies, our model provides a unifying framework for previously disjoint experimental observations. Finally, we propose a novel biomolecular controller that achieves near-perfect adaptation of cell-wide ribosome availability to changes in synthetic gene expression. Showcasing our model’s usefulness, we use it to determine the controller’s setpoint and operation range from its constituent genes’ parameters.

## 1 Introduction

The engineering of novel biological systems with useful applications, known as synthetic biology, holds the promise to address global challenges by revolutionising healthcare, agriculture, and manufacturing [1]. As for any engineered system, predictable behavior is a key requirement for synthetic biology designs. However, the forecasting of their performance is complicated by the interconnectedness of biological processes in living cells [2, 3, 4]. Indeed, even the gene circuit components that do not engage in direct biological interactions may exhibit interdependence caused by indirect couplings via shared cellular resources. Specifically, resource competition couplings arise when heterologous genes introduced into a host cell compete among themselves and with native genes for the same finite pools of resources that enable gene expression. Due to this competition, a synthetic gene circuit’s performance may not be reliably predicted based on the characterisation of its constituent modules in isolation [5, 6]. Moreover, the redirection of resources from native gene expression, known as gene expression burden, hinders cell growth and biomass accumulation [3, 4]. Resource couplings therefore present a key challenge in the engineering of bacterial, fungal, and mammalian cells [7, 8, 9].

A number of resource coupling mitigation strategies have been proposed [3]. For instance, synthetic gene expression can be maintained at levels that are low enough to have no significant effect on cellular homeostasis [10], whereas feedback mechanisms can improve a circuit’s robustness to perturbations such as resource couplings [7]. Moreover, synthetic genes can be expressed via orthogonal molecular machinery which is unused in native gene expression and forms a separate resource pool, thereby reducing crosstalk between engineered circuits and native processes [11]. Nonetheless, such crude approaches have limitations: while the low-expression strategy is unsuitable for applications requiring high protein production, the synthesis of orthogonal machinery or regulator proteins enabling feedback control can itself burden the cells [10]. Mathematical modeling of resource competition represents a promising approach to resource-aware design of biocircuits and the development of more sophisticated strategies to help mitigate resource competition couplings.

It is possible to explicitly model all stages of a substrate binding and unbinding the shared resource for which it competes with other substrate species. However, gene expression models can also be simplified to ensure better interpretability and lower computational complexity. The entire resource-dependent process can be described with an effective rate constant that reflects the concentrations of all competing substrates (Figure 1A), assuming that very fast association and dissociation make a resource-substrate complex’s concentration change very little on the time scale of other reactions in the cell [9, 12, 13]. Such simplified models facilitate biocircuit design by allowing users to easily determine a genetic module’s sensitivity to resource couplings [6] and to optimize design parameters like gene dosage and ribosome-binding site (RBS) strength, achieving desired outputs despite unwanted couplings [12, 14]. Furthermore, insights from modeling resource competition dynamics allow not only to develop more robust mechanisms for reducing the impact of unwanted couplings [7, 13], but also to leverage resource competition as a gene regulation mechanism to achieve other objectives, such as plasmid copy number-independent synthetic gene expression in mammalian cells [15].

**Figure 1:**
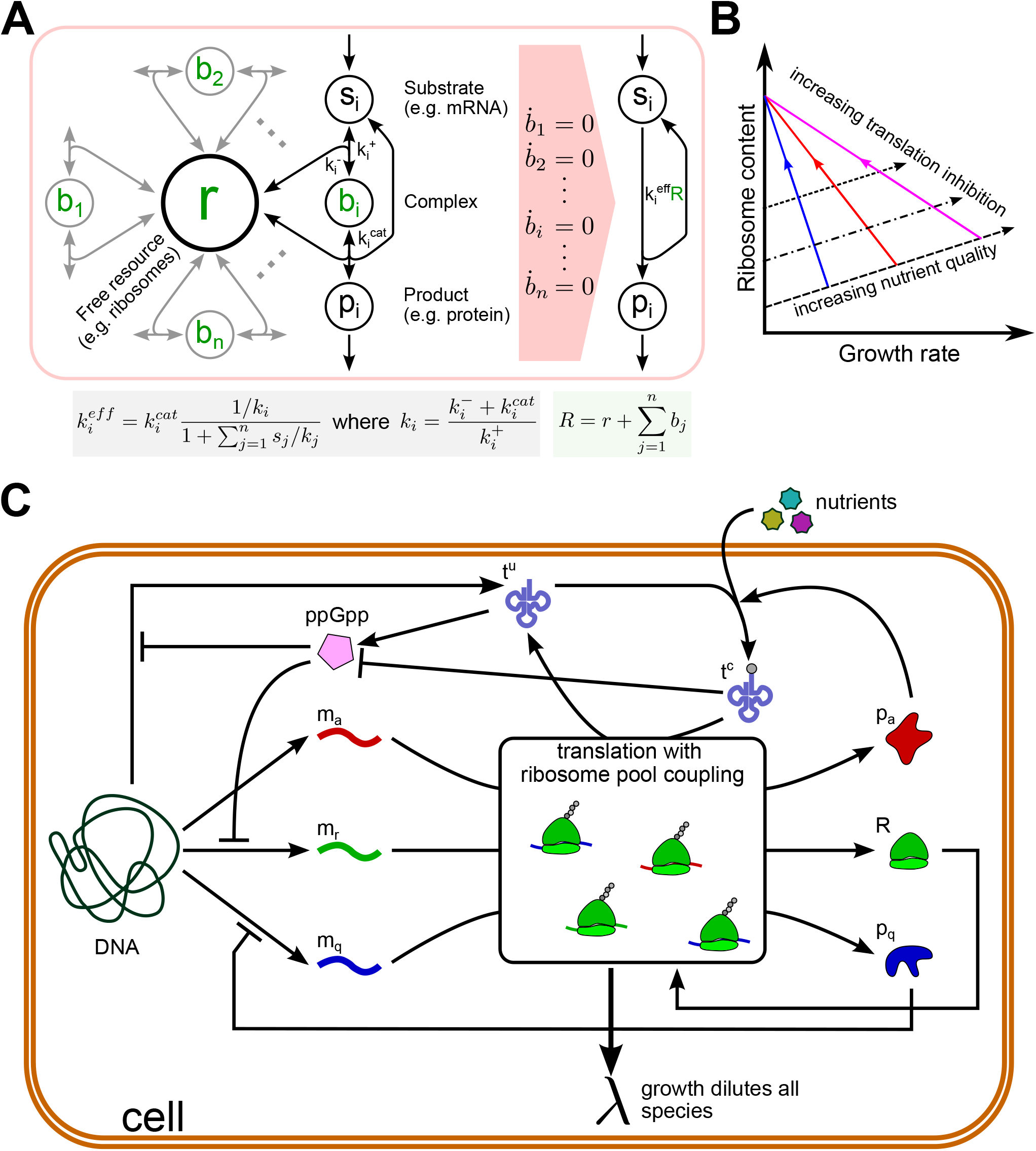
The proposed coarse-grained resource-aware bacterial cell model and its premises. **(A)** Instead of explicitly modeling the binding and unbinding of all competing substrates to the free resource (left), one can define the rate of a compound’s synthesis as a product of the total abundance of the resource *R*, which can be free (*r*) or bound to a substrate molecule (*b*_*i*_), and the effective rate constant 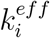(right). The effective rate constant depends on all competing species’ concentrations, affinities to the resource, and product synthesis rates [9]. **(B)** First (dashed line) and second (solid coloured lines) bacterial growth laws relate growth rate to ribosome content in different conditions. Formulated by Scott et al. [28], they postulate that the cell’s ribosome content increases linearly with the growth rate as the culture medium’s nutrient quality improves, but this relationship becomes inverse when translation is inhibited by an antibiotic. **(C)** Schematic of our host cell model. mRNAs are transcribed from genomic DNA and translated to make proteins. Competitive ribosome binding is not modeled explicitly – instead, effective rate constants relate each gene’s translation rate to the concentrations of protein precursors, ribosomes and all mRNAs in the cell. Ribosomes enable translation, whereas metabolic proteins catalyze nutrient import and tRNA aminoacylation. Native homeostatic regulation maintains housekeeping protein expression levels constant. The synthesis of tRNAs and ribosomal mRNAs is repressed by ppGpp, whose concentration reflects the reciprocal of the ratio of concentrations of charged and uncharged tRNA concentrations. The overall rate of protein synthesis defines the growth rate, since dilution due to growth must keep the cell’s total protein mass constant.

The analysis of resource couplings in bacteria is further complicated by the interplay between synthetic gene expression and cell growth rate. Since bacterial cells are fast-growing, dilution due to cell division is a major determinant of gene expression dynamics [16]. Accordingly, changes in the cell’s growth rate caused by gene expression burden can qualitatively change a gene circuit’s behavior [16, 17]. Moreover, the size of a bacterial cell’s pool of gene expression machinery is also related to its growth rate by the experimentally observed “bacterial growth laws” (Figure 1B) [18], which arise due to the cell optimizing its gene expression to maximize the steady-state growth rate in given environmental conditions [19, 20]. This limits the predictive power of current resource competition models, which rely on the assumption of constant growth rate and resource availability [6, 9, 10]. The development of bacterial resource-aware biocircuits thus calls for resource competition models that consider synthetic circuits within the context of the host cell and account for the impact of gene expression burden on cell growth.

Since allocating resources between different genes to achieve the highest growth rate can be considered an optimization problem, solving it allows to approximately predict gene expression in certain conditions [20, 21, 22]. However, ideal optimal control is not exhibited by actual living cells, which instead implement near-optimal gene regulation strategies via biological reactions [20]. Understanding these behaviors requires mechanistic cell models, which incorporate the trade-offs faced by living cells, such as the finiteness of the cell’s mass and its pool of gene expression machinery, energy, protein synthesis precursors, and other resources [23]. A range of such models with different levels of granularity exists, from whole-cell models [24] considering all known cellular processes to low-dimensional models, which group the cell’s genes with similar function and expression dynamics into just a few classes and only consider those cellular signalling pathways that are most relevant for the problem of resource allocation between the genes [21, 25]. In the investigation of the host cell’s interactions with synthetic gene circuits, capturing the cell’s growth regulation mechanisms and the various ways in which different design parameters of a synthetic device influence the cell must be balanced against maintaining minimal model complexity. Indeed, an overly detailed model may contain a high number of unknown or unidentifiable parameters [26], as well as hinder informative biocircuit analysis and complicate the understanding of core biochemical processes defining the cell’s state [27].

In this study, we propose a resource-aware coarse-grained mechanistic model of the bacterial cell that combines the present understanding of the host cell’s growth and metabolism regulation mechanisms with simplified resource competition analysis, enabled by the use of effective rate constants to capture the competitive binding of mRNAs to ribosomes. Our model therefore allows to easily derive analytical relations which can guide the choice of a biocircuit’s design parameters to achieve a desired behavior.

Parameterizing our model for *E. coli*, we show that it reproduces experimentally observed bacterial growth phenomena, as well as empirical relations between the burden-dependent reduction in growth rate and different quantities characterising heterologous gene expression in the cell. We also use our model to numerically reproduce the effect of resource competition on the activation dynamics of two bistable genetic switches hosted by the same cell. Finally, we showcase our model’s usefulness for the development of resource-aware biocircuits by leveraging it to propose and analyze a novel biomolecular controller for mitigating gene expression burden. By maintaining near-constant ribosome availability at a cell-wide level, it reduces the effect of indirect couplings via the shared ribosome pool.

## 2 Results

### 2.1 A coarse-grained mechanistic model capturing bacterial resource allocation and growth

We start by explaining the fundamental assumptions of our ordinary differential equation (ODE) model of the bacterial cell without any synthetic genes introduced to it, which is depicted in Figure 1C. We distinguish three classes of bacterial genes, respectively labelled *r, a*, and *q*: ribosomal, metabolic (catalysing tRNA aminoacylation), and housekeeping (whose expression stays constant regardless of culture conditions). Following a coarse-grained approach, all metabolic genes are treated as a single lumped gene; the same strategy is applied to the ribosomal genes. Meanwhile, the abundance of housekeeping proteins in the cell is assumed to be constant – namely, under a wide range of conditions, their share in the cell’s protein mass is fixed at 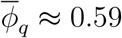 [21, 29]. Hence, we can avoid modeling their expression explicitly (Supplementary Information Section 1.4). Besides mRNA and protein expression, we model the concentrations of uncharged and charged (i.e., aminoacylated) tRNA molecules in the cell, since they play a key role in determining translation rates and resource allocation between gene classes [30]. Consequently, our mechanistic cell model is given by Equations (1)-(6), where *m*_*a*_ and *m*_*r*_ are respectively the concentrations of metabolic and ribosomal mRNAs, *p*_*a*_ is the metabolic protein concentration, *R* is the cell’s total ribosome count, and *t*^*c*^ and *t*^*u*^ are the concentrations of charged and uncharged tRNAs, respectively.

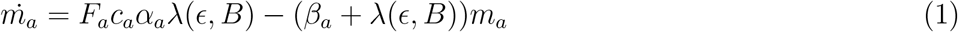

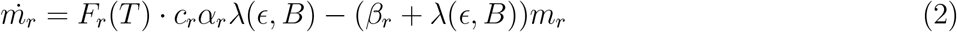

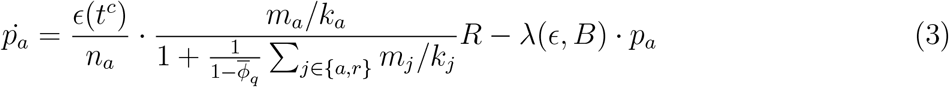

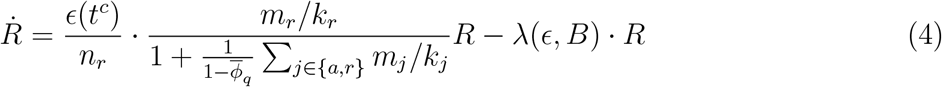

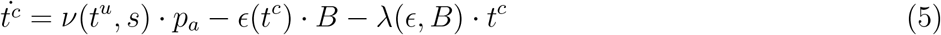

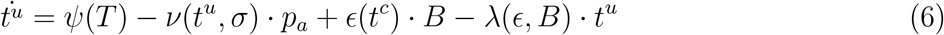

Gene expression and growth in bacteria are primarily affected by translational, rather than transcriptional, resource availability [6, 14, 31]. For simplicity, we therefore do not consider competition for transcriptional resources (e.g., RNA polymerases), so the rate of mRNA synthesis in Equations (1)-(2) is given by:

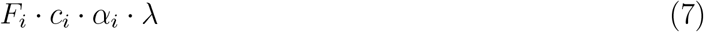

where *α*_*i*_ is the corresponding gene’s promoter strength and *c*_*i*_ is its DNA’s concentration, which for the cell’s native genes is assumed to be 1 *nM* as a convention (since the volume of an *E coli* cell is around 1 *μm*^3^, this is equivalent to one gene copy per cell [12]). *F*_*i*_ is the dimensionless transcription regulation function. For the ribosomal genes, *F*_*r*_ will be defined later in this section, while for the constitutively expressed metabolic genes [18, 28, 19] we use *F*_*a*_ ≡ 1. Finally, Equation (7) includes the cell’s growth rate *λ*, as mRNA production rate across the bacterial genome increases linearly with the growth rate [32]. Consequently, since the transcription rate is measured in *nM* of mRNA synthesized per hour and the units of *c*_*i*_ and *λ* are respectively *nM* and *h*^−1^, the promoter strength *α*_*i*_ is dimensionless.

To define protein synthesis rates in Equations (3)-(4), we adopt an approach similar to that used in [9] and [12]. Hence, in Supplementary Information Sections 1.2-1.3 we derive the effective translation rate constants:

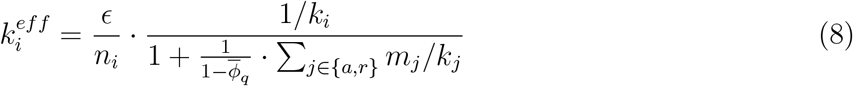

where

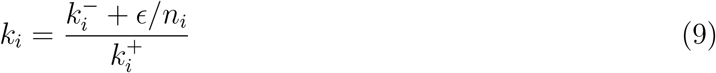

is the apparent mRNA-ribosome dissociation constant, determined by the binding and unbinding rates between the RBS and the ribosome (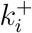 and 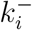, respectively) and the rate at which the ribosome completes translation to slide off the mRNA. The latter can be obtained by dividing *ϵ*, the translation elongation rate in amino acids per hour, by *n*_*i*_, the number of amino acid residues in the protein encoded by the gene.

As for the removal of molecules, all species are diluted at a rate *λ* due to the cell growing and dividing. While the overwhelming majority of proteins is not actively degraded in the bacterial cell [33], mRNA degradation, conversely, is widespread and happens at a high rate [34]. Hence, we include additional constant degradation terms *β*_*a*_ and *β*_*r*_ in Equations (1)-(2), which describe mRNA concentration dynamics.

The cell growth rate *λ* is related to the cell’s state by the “finite proteome cellular trade-off”, identified by Weisse et al. [27]. While the shares of different protein classes in the cell’s protein mass can vary, the total mass of proteins per unit of cell volume has been observed to be constant in the exponential growth phase [35]. Consequently, the overall rate of production of all proteins must equal the total rate of protein mass dilution due to cell division. If *M* is defined as the mass of proteins (in amino acid residues) in the average volume of the cell over the cell cycle and

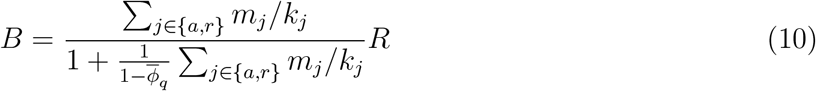

is the total concentration of actively translating ribosomes (Supplementary Information Section 1.3), Equation S8 in the Supplementary Information shows that *λ* can be found as:

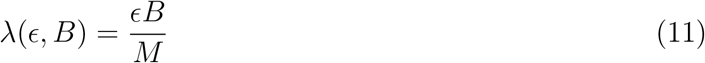

We now proceed to define the translation elongation rate *ϵ*, which depends on *t*^*c*^, the size of the pool of protein synthesis precursors (i.e., charged tRNAs) [19]. This relationship can be described with Michaelis-Menten kinetics [20] as shown in Equation (12), where *ϵ*_*max*_ is the maximum possible translation elongation rate and *K*_*ϵ*_ is the half-saturation constant.

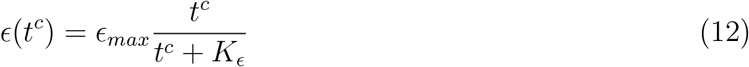

To replenish the protein precursors consumed during translation, metabolic proteins aminoacylate the uncharged tRNAs, *t*^*u*^. Once again adopting a Michaelis-Menten approach [21], we can describe the rate of tRNA charging using Equation (13).

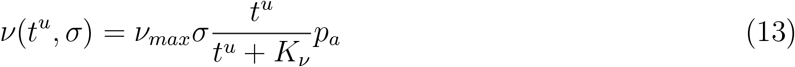

Here, 0 *< σ* ≤ 1 is a dimensionless constant quantifying the growth substrate’s nutrient quality, *K*_*ν*_ is the half-saturation constant, and *ν*_*max*_ is the maximum tRNA aminoacylation rate per metabolic enzyme.

Finally, we consider how the cell allocates resources between the expression of different genes, which is captured by the ribosomal gene transcription regulation function *F*_*r*_. The near-optimal regulation of bacterial gene expression, aimed at maximizing the steady-state cell growth rate, is believed to be enabled by the guanosine tetraphosphate (ppGpp) signalling pathway [19, 20]. We implement it in our model according to the recently proposed Flux-Parity Regulation theory [21], which states that the cell tries to equate and maximize its protein synthesis and tRNA aminoacylation fluxes. A growing body of evidence suggests that ppGpp concentration ([*ppGpp*]) reflects the ratio of the aminoacylated and uncharged tRNA concentrations [30, 36]. Consequently, the cellular concentration of ppGpp is described by the variable *T*, where

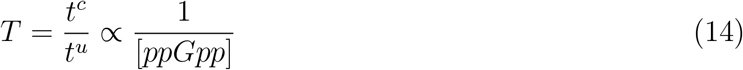

Notably, ppGpp represses ribosome synthesis [37], which can be captured by Hill kinetics outlined in Equation (15), where *τ* is the half-saturation constant.

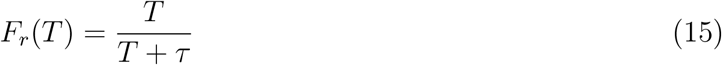

Since tRNA genes are co-regulated with the ribosomal genes [38], tRNA production can likewise be described by Equation (16), with *ψ*_*max*_ being the maximum tRNA transcription rate.

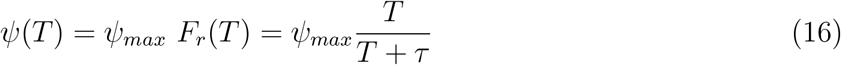

### 2.2 The proposed cell model predicts growth phenomena in *E. coli*

In order to validate model predictions against experimental data, we parameterized it for *E. coli*, which is the most studied bacterial model organism [18, 21, 30, 39] and one of the most popular host organisms in synthetic biology [40]. Most parameter values, displayed in Supplementary Table S1, were taken from literature. The rest were fitted to experimental measurements of growth rate and ribosomal mass fraction of *E. coli* subjected to different concentrations of the translation-inhibiting antibiotic chloramphenicol in various growth media [18, 21] as described in Methods and Supplementary Information Section 2.

Our model’s steady-state predictions for different culture conditions are presented in Figure 2A. As can be seen, they are generally consistent with empirically proposed bacterial growth laws illustrated in Figure 1B [18, 28]. Indeed, when nutrient quality improves and chloramphenicol levels remain unchanged, the cell’s ribosome content increases linearly with the growth rate (dashed lines), obeying the first bacterial growth law. Moreover, augmenting translation elongation inhibition for the same nutrient quality produces an inverse proportionality between the ribosome content and growth rate for chloramphenicol levels of up to 4*μM* (solid lines). We also confirmed the consistency of our model predictions for the cell’s ribosome content, translation elongation rate, and ppGpp level with experimental results. To this end, we varied the nutrient quality factor *σ* without inhibiting translation and compared our model’s predictions with experimental data from the 37 different experimental studies compiled by Chure and Cremer [21] (Figure 2B–D).

**Figure 2:**
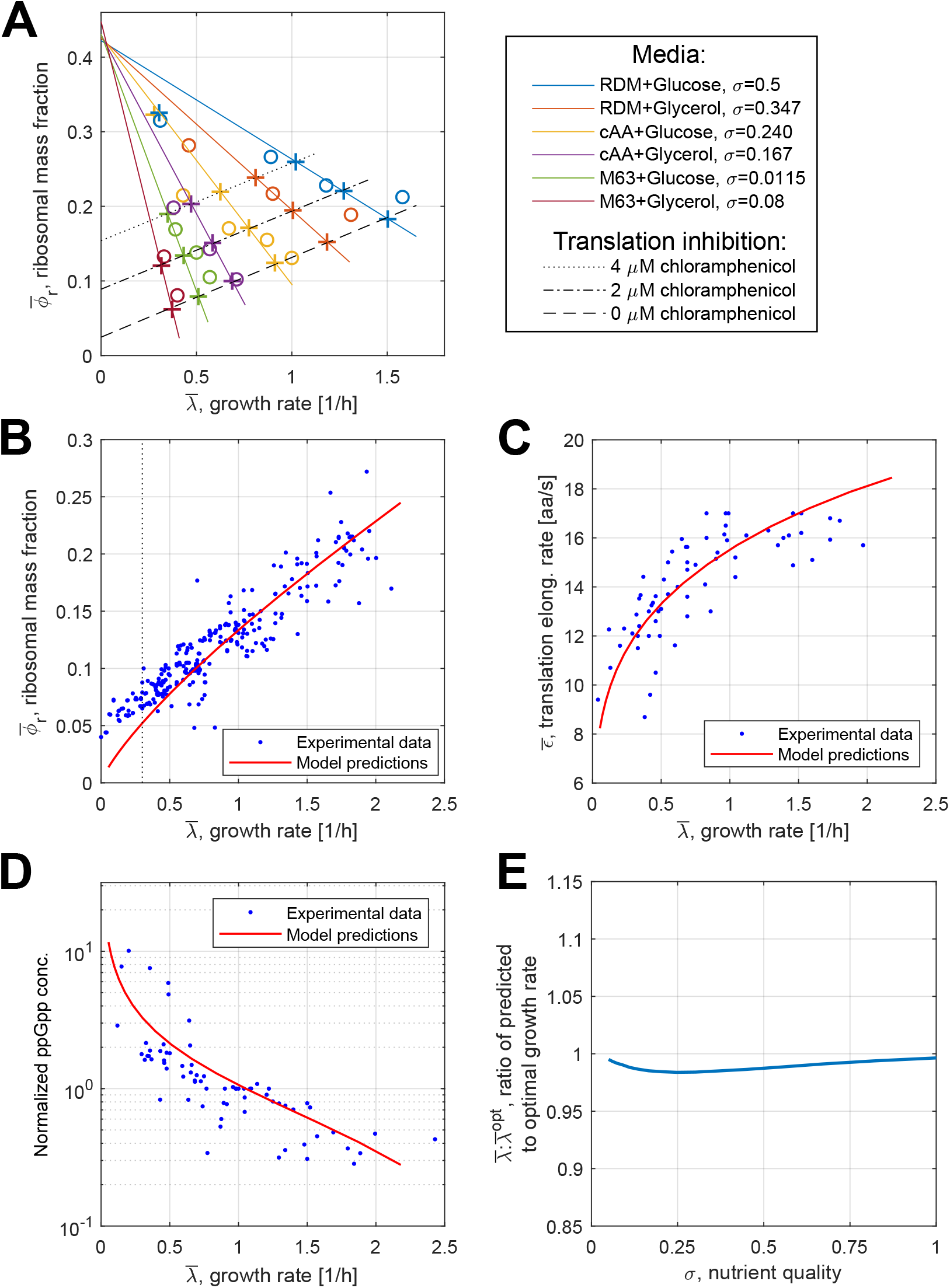
Growth phenomena predicted by our model. **(A)** Experimental measurements by Scott et al. [18] used in parameter fitting (circles) and model predictions (crosses) for steady-state ribosomal mass fractions and growth rates. The first and the second growth laws are illustrated by dashed and solid lines, respectively. Note that the growth law fits do not include the two predicted points for translation inhibition with 8 *μM* of chloramphenicol, situated in the plot’s top left corner. **(B, C, D)** Comparison of model predictions for the cell in steady state with experimental data from previous studies [21]. In (B), the dotted vertical line denotes the 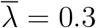 threshold, left of which model predictions for 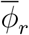 significantly diverge from the experimental data. In (D), the ppGpp levels are normalized to the reference value for which 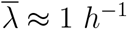. **(E)** Ratio of the steady-state growth rate 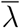 predicted by our model to the optimal growth rate 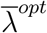 for *σ* varied from 0.05 to 1.

While Figure 2B supports the linear-like bacterial growth law dependence between ribosome content and growth rate, for growth rates below *λ* ≈ 0.3 *h*^−1^ our model predictions take a nonlinear downturn, diverging from experimental measurements. Moreover, for *E. coli* subjected to 8*μM* of chloramphenicol, predictions are situated off the growth law trendlines. Notably, these discordances arise in highly unfavorable conditions – that is, strong inhibition of translation or very poor nutrient quality of the medium. Hence, to some extent they may be attributed to experimental errors, since it is difficult to ensure that measurements in such conditions pertain to cells in steady state [21]. However, hostile environments can also trigger the cell’s stress response mechanisms unconsidered by our model, such as ribosome inactivation, preserving the bacterium’s ability to synthesize proteins in adverse conditions [39]. Furthermore, since mRNA synthesis rate depends on the cell’s growth rate [32], low nutrient culture media reduce the overall concentration of transcripts in the cell. This leads to mRNA scarcity, rather than translational resource allocation that our model focuses on, becoming the key determinant of the growth rate [22]. Together, these factors limit our model’s predictive power in unfavorable conditions.

Finally, we confirmed whether ppGpp regulation in our model achieves near-optimal steadystate growth rates in a wide range of culturing conditions. For each medium nutrient quality *σ* considered, we did the following. First, we recorded the steady-state cell growth rate 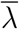 (here, the bar notation indicates a variable’s steady-state value) predicted by our model. According to it, *T*, the inverse of ppGpp concentration, reflects the charged and uncharged tRNA levels as detailed by Equation (14). Then, we assumed that ppGpp concentration in the cell is instead constant, ran simulations for different fixed values of *T* and identified the maximum steady-state cell growth rate 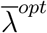 across all considered values. As shown in Figure 2D, for 0.05 ≤ *σ* ≤ 1 the growth rate forecast by our ppGpp regulation model was always very close (within 2%) to the optimal value, which supports the notion of near-optimal control of resource allocation.

### 2.3 Including Heterologous Gene Expression in Our Cell Model

Our cell model can be extended to consider heterologous gene expression. This allows to simulate gene circuit dynamics while accounting for resource couplings and the effects of heterologous gene expression on the host’s metabolism that can influence the biocircuit’s performance, such as growth rate changes. In addition to native genes, let there be a set of *L* heterologous genes *X* = {*x*_1_, …, *x*_*L*_}. To model their expression, we add to Equations (1)-(6) a pair of ODEs for each heterologous gene, which describes its transcription and translation (Supplementary Information Section 3). For a synthetic gene *x*_*l*_, they are:

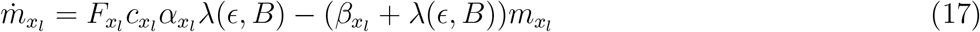

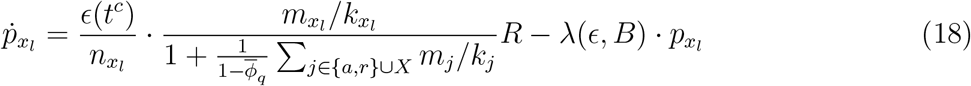

where all parameters have a similar definition to those of the parameters describing the native genes, while the transcription regulation function 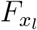 is circuit- and gene-specific. Likewise to the original host cell model, we assume that transcriptional resource couplings are insignificant. Meanwhile, the constant housekeeping protein mass fraction 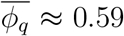 remains the same despite synthetic gene expression [18, 28, 29].

Consequently, the model’s original ODEs for native mRNA and tRNA levels are not aletered by the addition of synthetic genes. Conversely, Equations (3) and (4) describe protein concentrations and thus include the terms for translation. Therefore, they must be amended to consider additional synthetic mRNAs competing for ribosomes:

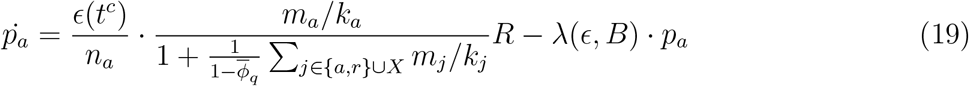

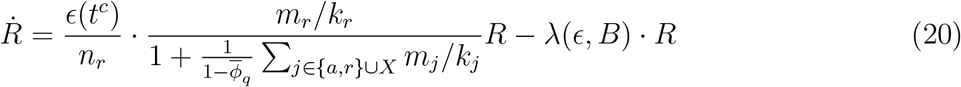

To demonstrate how our model captures the effects of resource competition on synthetic gene expression, we used it to recreate the “winner-takes-all” phenomenon, a known case of biocircuit behavior being altered by resource couplings [5]. Consider a gene that encodes a protein which, upon being allosterically modulated by an inducer molecule, acts as a transcription factor to activate its own expression. In isolation, this system can act as a bistable switch, since the concentration of the protein has two stable equilibria – one with a low expression rate and one involving high expression levels. Thus, the system converges to one of the equilibria, depending on whose basin of attraction its initial condition belongs to. These equilibria and the border between their basins of attraction can be shifted by varying the inducer’s concentration in the medium [41]. In a “winner-takes-all” scenario, two such switches are found in the same cell, interacting indirectly via the shared resource pool (Figure 3A). Therefore, if one switch (“the winner”) reaches its high-expression equilibrium before the other switch, the winner’s abundant transcripts outcompete the other switch gene’s mRNAs. Consequently, with the second protein’s production hindered by reduced ribosome availability, its concentration is prevented from reaching the high-expression equilibrium value. Meanwhile, the winner switch gene uses up almost all of the expression resources [5].

**Figure 3:**
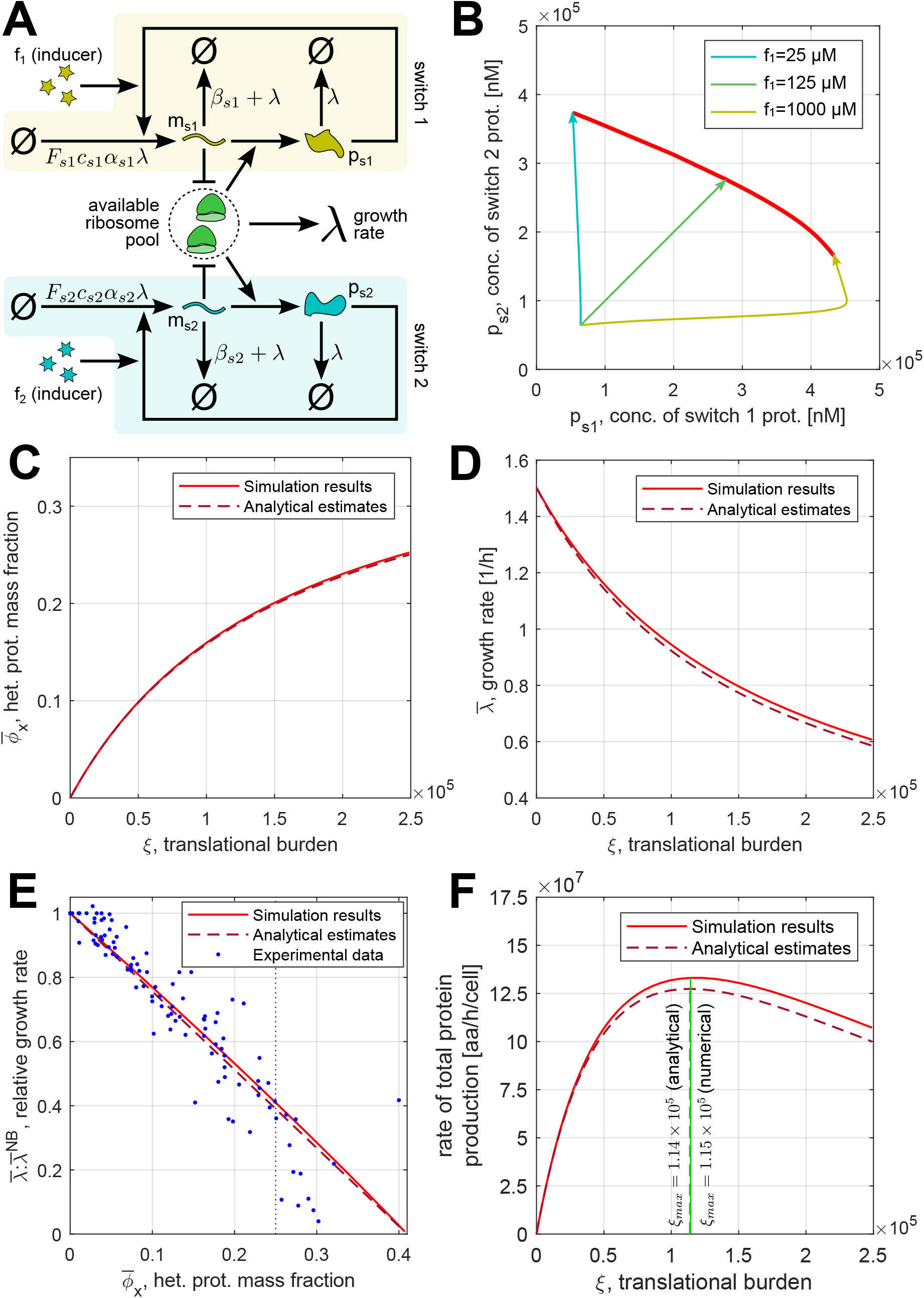
Numerical and analytical predictions of heterologous gene expression and its effects on the host cell. **(A)** Two bistable switches in the same cell interact via the shared resource pool, which may lead to “winner-takes-all” scenarios where the activation of one switch suppresses the other [5]. **(B)** Phase plane diagram showing the two-switch system’s behavior upon simultaneous addition of inducer 1 and inducer 2 to the culture medium. Thin lines with arrowheads: the system trajectories for different concentrations of inducer 1 being added. Bold red line: system’s final steady states as the concentration of inducer 1 varies between 25 *μM* and 1000 *μM*. In all cases, the concentration of inducer 2 being added is 125 *μM*. **(C, D)** Hill activation and repression functions respectively relate translational burden to the steady-state heterologous protein mass fraction (C) and cell growth rate (D). **(E)** In the steady state, 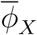, the mass fraction of heterologous proteins in the cell, is linearly related to 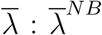, the ratio of the corresponding growth rate to that without burden. The dotted vertical line denotes the 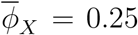 threshold, right of which model predictions become unreliable. **(F)** The rate of total protein production per cell as a function of the translational burden. *ξ*_*max*_ (green lines) is the burden value maximizing heterologous protein synthesis. All simulations assume *σ* = 0.5; the values of other parameters can be found in Supplementary Tables S4 and S5.

We modeled the system of two bistable switch genes *s*_1_ and *s*_2_ by describing the corresponding mRNA and protein concentrations with Equations S79-S82 in the Supplementary Information, based on Equations (17)-(18). The system was parameterized as outlined in Supplementary Table S5. Denoting the concentration of gene *s*_*i*_’s inducer molecule as *f*_*i*_, we define its gene transcription regulation function as follows:

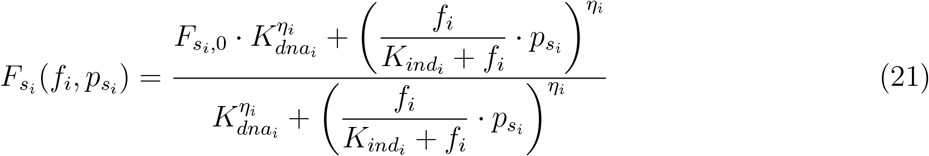

In Equation (21), the allosteric modulation of the protein by inducer binding is described with a Hill function, where 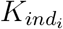 is the dissociation constant between the inducer and the protein target [12]. For the binding of the inducer-protein complex to the promoter DNA, Hill kinetics are assumed, with 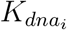 being the half-saturation constant and *η*_*i*_ the corresponding Hill coefficient, which captures the cooperativity of binding [41]. 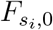 represents the gene expression regulation function’s value in the absence of any transcription activation factors due to leaky expression.

Figure 3B displays the phase plane plot for the two synthetic proteins’ concentrations when different amounts of each inducer are added to the medium. The phase plane trajectories demonstrate that when a high inducer concentration causes one switch to reach the high-expression equilibrium faster than the other, the latter switch’s activation is blocked. This is shown by the blue and yellow arrows in Figure 3B, standing respectively for 25 *μM* = *f*_1_ *< f*_2_ = 125 *μM* and 1000 *μM* = *f*_1_ *> f*_2_ = 125 *μM*. Equal induction of both switches leads to them being simultaneously activated (green arrow in Figure 3B for *f*_1_ = *f*_2_ = 125 *μM*). These simulation results are in line with the experimental observations for such two-switch systems in bacteria, as well as the current understanding of the mechanisms underlying “winner-takes-all” phenotypes [5], which supports our model’s applicability to the study of changes in synthetic circuits’ performance caused by resource competition couplings. However, if even in the activated state the expression of a switch gene is low, its contribution to resource competition in the cell may be insufficient to affect the other switch and produce “winner-takes-all” behavior (Supplementary Figure S3). Using our model to simulate a gene circuit’s dynamics with given design parameters could therefore allow to determine whether it exhibits resource competition phenomena of interest.

### 2.4 Analytical Predictions Reveal How Heterologous Gene Expression Affects the Cell

Besides enabling numerical prediction of circuit behavior, with some simplifying assumptions our model allows to derive analytical relations that reveal the effect of different parameters on the steady-state values of different variables. As we demonstrate in Supplementary Figure S2, simulations show that in nutrient-rich media (i.e., for *σ* ≥ 0.39) the translation elongation rate and ribosomal gene transcription regulation change very little over a wide range of heterologous gene expression rates. This allows us to assume that 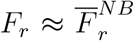 and 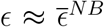 (hence also 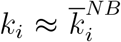), regardless of which synthetic genes are present. The ^*NB*^ (“no burden”) index denotes the steadystate values in absence of any synthetic gene expression – that is, *X* = ∅. These steady-state values can be found by simulating the host cell model without synthetic genes for a given *σ*. Additionally, we assume that all mRNA molecules in the system decay at roughly the same rate, i.e., *β*_*i*_ ≈ *β*_*j*_, ∀*i, j* ∈ {*a, j*} ∪ *X*. Importantly, while the transcripts of individual native *E. coli* genes have very different degradation rates [34], the coarse-grained nature of our model means that this assumption only concerns the *average* degradation rates across the native gene classes, each of them spanning many genes.

The “translational burden” imposed on the cell by the synthetic genes *x*_1_, …, *x*_*L*_ ∈ *X* – that is, the additional competition for ribosomes experienced by native genes – can be quantified by a lumped factor *ξ* as shown in Equation (22), where 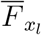 is the steady-state value of gene *x*_*l*_’s transcription regulation function.

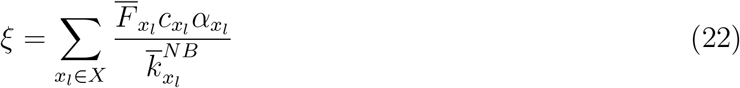

The steady-state mass fraction of heterologous protein in the cell as a function of *ξ* can then be estimated as shown in Equation (23), while the dependence of the steady-state cell growth rate on burden is captured by Equation (24) (see Supplementary Information Section 3.2 for the derivations).

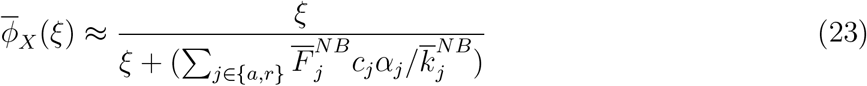

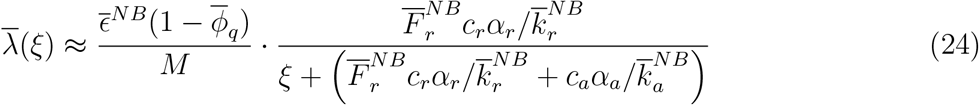

Furthermore, the change in the cell growth rate relative to 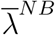 – that is, 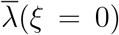 – can be related to the total mass fraction of all heterologous proteins in the cell 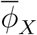 (Supplementary Information Section 3.3):

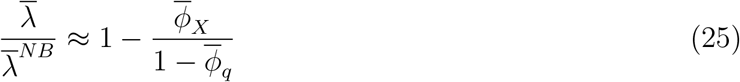

Several different empirical relations, such as Hill or linear dependencies, have been suggested to link gene expression burden to the reduction in cell growth rate [6, 18]. Therefore, previous works have considered different formulae as part of their gene expression models in order to ensure that the modeling outcomes stay valid regardless of the assumed burden-growth dependency [42]. However, Equations (24) and (25) hint that these relations may not be mutually exclusive, but rather apply to different quantities describing heterologous gene expression. Namely, Equation (24) relates growth rate to the translational burden – that is, a measure of the synthetic genes’ resource demand, determined by the circuit’s design parameters – giving rise to a Hill relation akin to that used in the work of McBride et al. [6]. If growth rate is linked to the heterologous protein yield, which is affected by resource couplings, a linear dependency from Equation (25) emerges, matching the observations of Scott et al. [18]. Our model therefore provides a possible unifying framework to explain different empirical laws describing the dependence of cell growth rates on synthetic gene expression.

To compare our analytical estimates with the model’s numerical predictions for different extents of translational burden experienced by the host, we simulated the expression of a constitutive heterologous gene by an *E. coli* cell (parameterized in Supplementary Table S4). The translational burden *ξ* exerted on the host by the synthetic gene was varied by sweeping through different values of the gene’s DNA concentration from 1 *nM*. The results are plotted in Figure 3C-E. As can be observed, our approximate analytical predictions of Equation (23), (24), and (25) closely follow the steady-state values obtained by numerical simulation (Figure 3D, E, and F, respectively). In Figure 3C, we can also see that both the analytical and numerical predictions are largely concordant with experimental data compiled by Chure and Cremer [21], although the real and predicted measurements diverge when the heterologous protein overexpression stress is very high 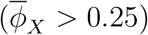. The explanation for this discrepancy is likely similar to the general reasons why our model’s predictions do not match experimental data in highly unfavorable conditions. Outlined in Results Section 2.2, they include measurement errors [21], ribosome inactivation [39], and the dependence of slowly dividing cells’ growth rate on their total mRNA content [22].

Analytical relations derived using our model can be useful in the design of heterologous gene expression systems. As an example, consider a population of *N E. coli* cells expressing a constitutive heterologous protein of interest *p*_*poi*_ and dying at a constant rate *δ*, modeled by Equation (26):

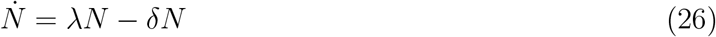

We can analytically find the optimal value of translational burden *ξ*_*max*_ that maximizes the production rate *μ* of the protein of interest by the cell population (Supplementary Information Section 3.4). This allows to derive an analytical optimality condition based on the gene of interest’s design parameters, namely its DNA copy number *c*_*poi*_, promoter strength *α*_*poi*_, and apparent mRNA-ribosome dissociation constant 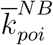 (reflecting the RBS strength):

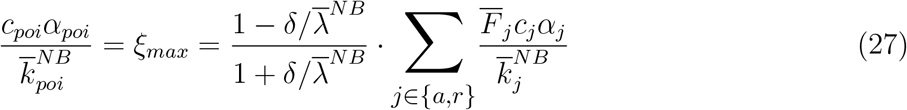

In Figure 3F, we plot the total protein production rate (calculated according to Equation S75 in the Supplementary Information) as a function of the gene expression burden *ξ* assuming that *δ* = 0.25 *h*^−1^. Notably, *ξ*_*max*_ yielded by Equation (27) lies within 1.03% of the numerically found optimal value.

### 2.5 A proportional-integral biomolecular controller for mitigating gene expression burden

To demonstrate how our model can aid the development of resource-aware circuits, we used it to design and analyze a biomolecular controller that mitigates indirect resource couplings via the shared ribosome pool, reducing the impact of gene expression burden. Usually, robustness to resource couplings is achieved for a single variable by applying feedback or, alternatively, for the limited set of synthetic genes that draw resources from an orthogonal pool, whose size is regulated by a biomolecular controller [7]. Conversely, the design we propose here aims to reduce fluctuations in resource availability at the whole-cell level. Our cell model is particularly suited for studying such controllers: by considering the expression of not just synthetic but all genes in the cell, it captures cell-wide resource couplings that our controller seeks to minimise. Meanwhile, the analysis of these couplings is facilitated by the cell model’s effective rate constant framework.

The proposed circuit, whose schematic is given in Figure 4, employs the Antithetic Integral Feedback (AIF) motif, which can be used to maintain a physiological variable of interest at a desired setpoint value [43]. It comprises an annihilator and an actuator species, where the annihilator’s synthesis rate depends on the controlled variable and the actuator species is produced at a constant rate, which sets the desired reference value. Since one annihilator and one actuator molecule can react to disable each other, the concentration of remaining non-annihilated actuator molecules reflects the integral of the error between the controlled variable and the reference. By influencing the rest of the system, the actuator molecules ensure robust perfect adaptation (RPA) to disturbances [43]. To control our variable of interest – that is, ribosome availability in the cell – we implement this motif using RNA logic, where the actuator is a protein-encoding mRNA and the annihilator is a small RNA (sRNA) that can bind it to form a rapidly degraded complex [44, 45]. As it does not involve protein synthesis, this RNA-based implementation is neither affected nor disrupted by translational resource couplings that we seek to mitigate, whereas transcriptional couplings in bacteria are largely insignificant [6, 14, 31].

**Figure 4:**
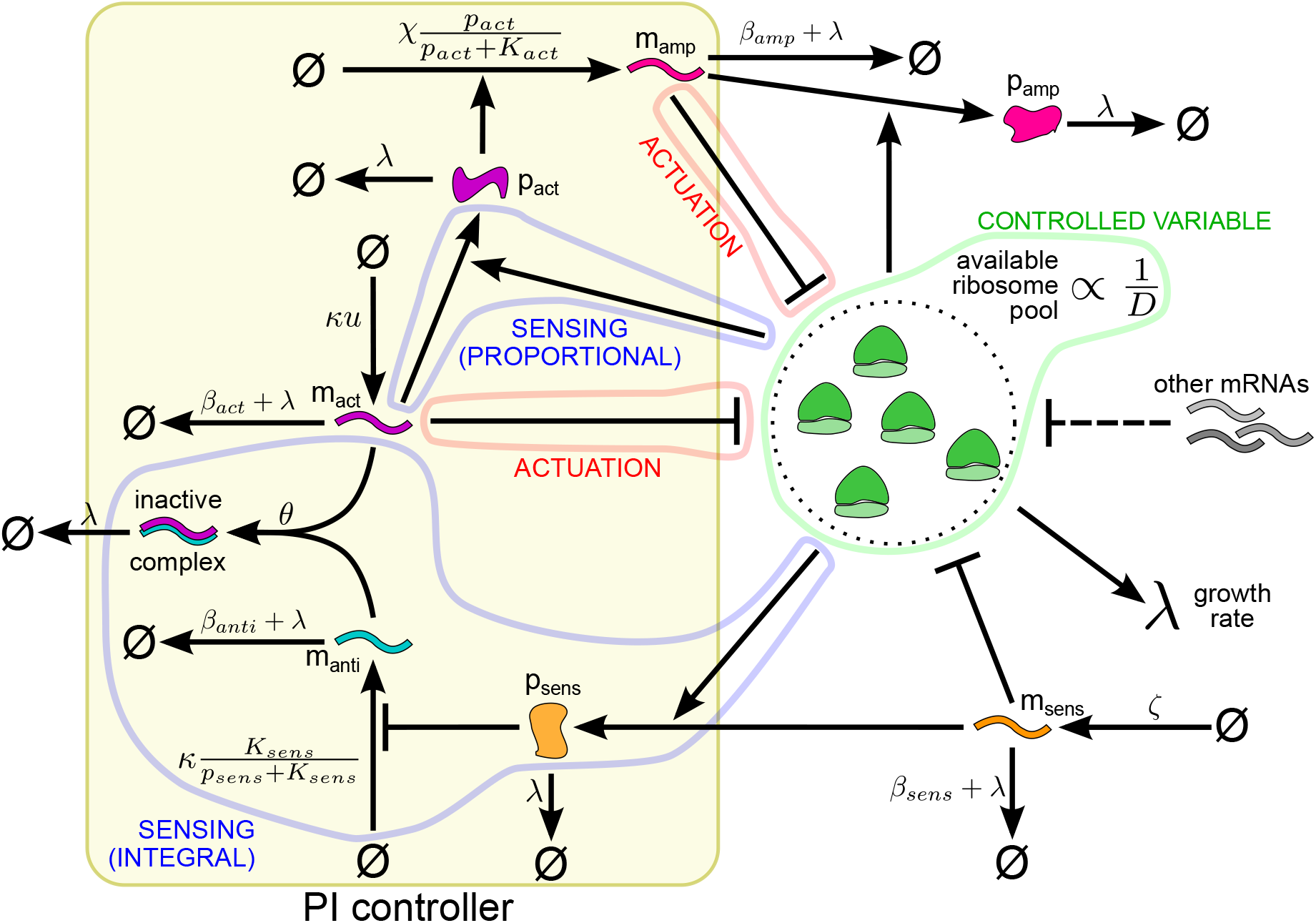
Schematic of the proportional-integral biomolecular controller mitigating gene expression burden. If some disturbance increases the number of mRNAs competing for ribosomes, the concentration of the constitutive protein *p*_*sens*_ falls, which activates the production of annihilator RNA *m*_*anti*_. Hence, the concentration of the actuator mRNA *m*_*act*_ decreases; so does the expression of the amplifier mRNA *m*_*amp*_, regulated by the actuator protein *p*_*act*_. This brings competition for ribosomes down to the original level. Integral feedback is enabled by the AIF motif involving *m*_*act*_ and *m*_*anti*_, while the proportional component arises from *p*_*act*_ expression being affected by ribosome availability.

The circuit, illustrated in Figure 4 and modeled by Equations S86-S92 in the Supplementary Information, comprises four genes: the “sensor” *sens*, the annihilator *anti*, the actuator *act*, and the “amplifier” *amp*. Ribosomal competition affects the level of the constitutively expressed transcription factor that is encoded by the sensor gene and regulates the annihilator sRNA’s transcription. Actuation happens by changing the concentrations of the transcripts competing for ribosomes. While the translation of the actuator mRNA sequesters some ribosomes, the protein encoded by it also regulates the transcription of an additional amplifier gene. This amplification is needed because heterologous mRNA concentration can significantly affect the ribosomal competition landscape only if it is large enough to be comparable to the abundance of native transcripts. However, a substantial amount of actuator mRNAs is sequestered by the annihilator. Hence, achieving such a large concentration of heterologous transcripts by only expressing the actuator gene would require very high transcription rates. We therefore circumvent this issue by tasking another mRNA, which is not annihilated by any sRNA, with implementing the bulk of the control action.

The variable of interest, whose fluctuations our controller aims to minimise, is the availability of ribosomes in the cell. It can be quantified by the “resource competition denominator”

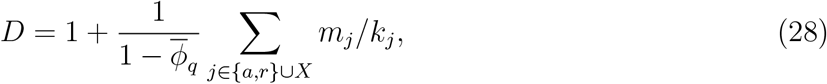

called so because in Equations (18)-(20) it is the denominator of the effective protein synthesis rate constant, which adjusts the corresponding translation rate in view of competition from other transcripts. However, the controlled variable *D* cannot be directly sensed by the RNA-based AIF motif, hence the need for a constitutive sensor protein whose levels depend on resource competition.

Therefore, the control error 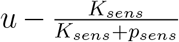 perceived by the motif is proportional to the difference between the actuator and the annihilator RNAs’ synthesis rates, which respectively equal *κu* and 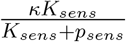 (Figure 4). Meanwhile, since the actuator and amplifier genes belong to the set of all heterologous genes *X* in Equation (28), the control action is exerted directly on *D*. Additionally, the actuator protein’s synthesis rate is itself susceptible to resource competition, and therefore is a multiple of ^1^/_*D*_. Therefore, the control action exerted through *m*_*amp*_ (whose synthesis is regulated by *p*_*act*_) has a multiplicative proportional component besides integral action [46].

While the controlled variable is the resource competition denominator *D*, the sensed error is a function of the sensor protein’s concentration *p*_*s*_ and the reference value *u*, which characterizes actuator mRNA production. In order to tune the setpoint value of *D* for a particular application of interest, it is therefore necessary to understand how it is related to *u* via *p*_*s*_. Knowing that integral feedback reduces the steady-state control error to zero, we can retrieve the steady-state setpoint value of *D* as a function of the controller’s parameters according to Equation (29) (Supplementary Information Section 4.2.2). The meaning of the circuit’s parameters found in this and other equations regarding our proportional-integral controller is illustrated in Figure 4 and explained in Supplementary Tables S7-S8.

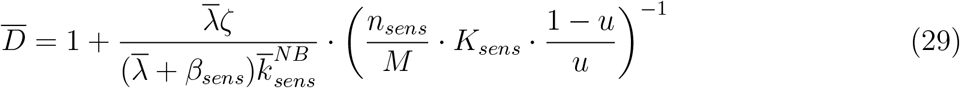

Likewise, it is possible to find the steady-state growth rate maintained by the controller:

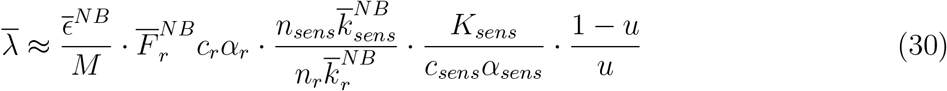

Moreover, we can analytically estimate the controller’s operation range – that is, the range of disturbance magnitudes which our design can negate to maintain a near-constant ribosome availability (Supplementary Information Section 4.2.2). If the cell is endowed with *L* synthetic genes {*x*_1_, *x*_2_, … *x*_*L*_} besides those of the proportional-integral controller, the disturbance caused by expressing them is mitigable by our controller only if:

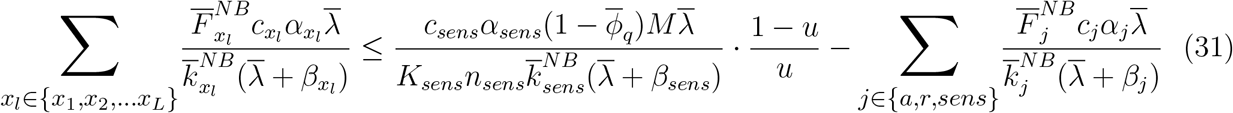

In order to test the proposed controller’s performance and the accuracy of our estimates, we used our cell model to simulate how the proportional-integral controller reacts to the appearance of an additional mRNA species competing for the shared pool of ribosomes (Figure 5A) and plotted the outcome in Figure 5B–F. Figure 5B–C shows that the amplifier mRNA levels fall in response to a step disturbance so as to restore the original extent of resource competition. Consequently, the adaptation error – that is, the difference between the variables’ values before and after disturbance – amounted to only 1.15% and 0.15% for the cell’s growth rate and the resource competition denominator, respectively. The observed non-zero adaptation errors can be explained by “leakiness”.

**Figure 5:**
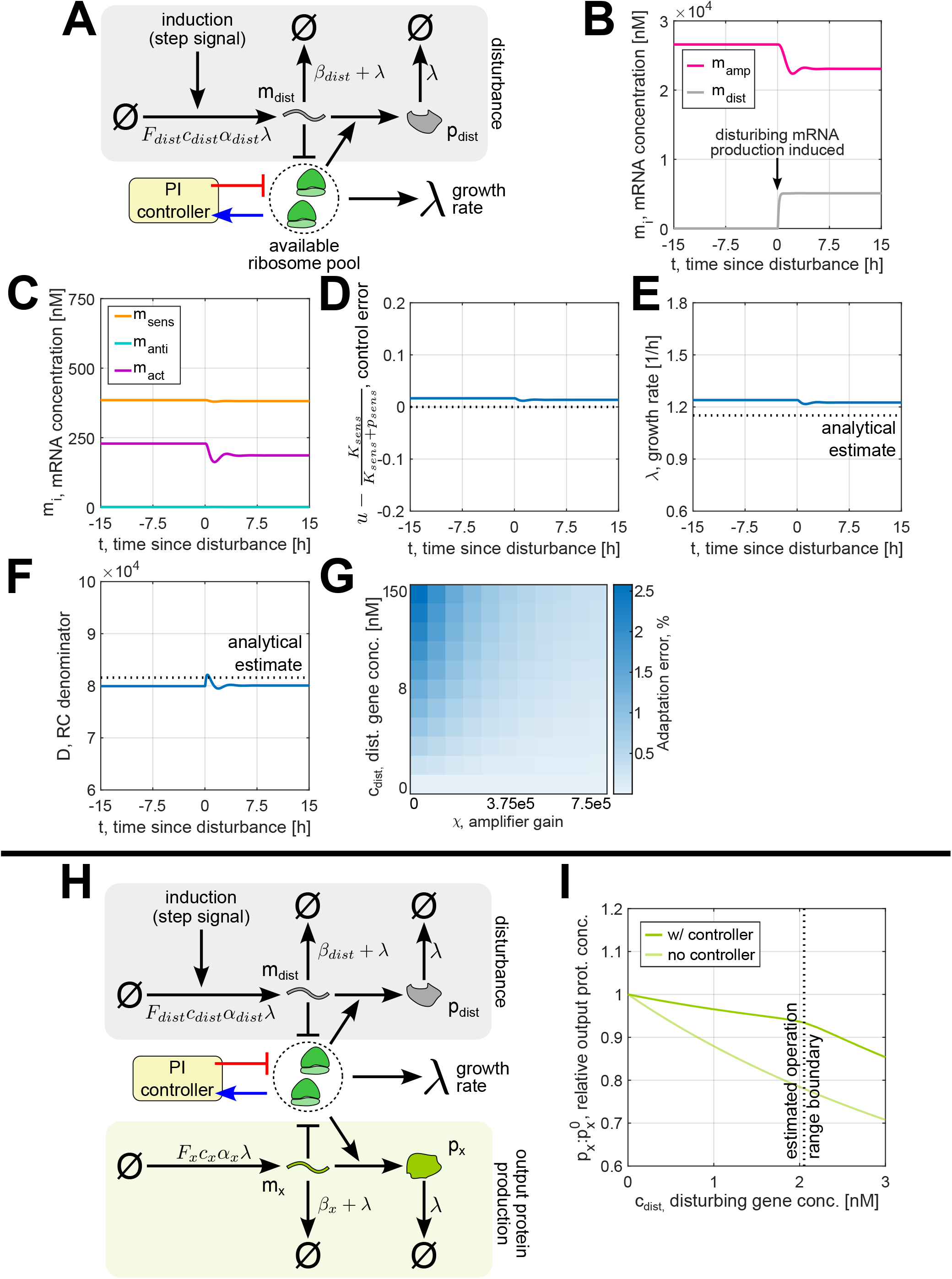
Simulation of the proportional-integral controller’s response to disturbance. **(A)** First case: the expression of an additional disturbing synthetic gene *dist* is induced, so the controller counteracts the resulting change in burden. **(B-F)** Example simulation of the first case with the parameters given in Supplementary Table S7. Figures (B) and (C) display the evolution of mRNA concentrations over time, while figures (D), (E), (F) show the control error sensed by the AIF motif, the cell’s growth rate, and the resource competition denominator *D*, which is the circuit’s controlled variable. **(G)** The controller’s adaptation error, i.e., the difference between the steady-state value of *D* before and after disturbance, for various values of the amplifier gain *χ* and the disturbing gene’s DNA concentration *c*_*dist*_. All other circuit parameters are taken from Supplementary Table S7. **(H)** Second case: besides the disturbing gene, the cell also contains the output gene of interest *x*, whose protein concentration the controller aims to keep constant despite the disturbance. **(G)** Ratio of the output protein’s level *p*_*x*_ to its concentration 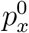 in absence of disturbance, plotted against the total production rate of the disturbing mRNA in the cell. All simulation parameters apart from *c*_*dist*_, which is varied, can be found in Supplementary Table S7.

This phenomenon arises in AIF controllers when the rate of the actuator’s and annihilator’s degradation and dilution is non-negligible compared to their rate of mutual elimination, preventing the output species from achieving RPA. The steady-state control error was also found to be non-zero, comprising 3.33% of the reference *u* (Figure 5D). Meanwhile, the analytical estimates for *λ* and *D* were found to be within 7.69% and 2.0% from the numerically determined values (Figure 5E–F). Besides the assumptions of constant *ϵ* and *F*_*r*_ made to retrieve the analytical predictions (see Results Section 2.4), leakiness has also contributed to these discrepancies, since it was neglected in the derivation of Equations (29)-(31) (Supplementary Information Section 4.2.2).

While leakiness can arbitrarily deteriorate the performance of AIF controllers, in our case the controlled variable *D* and the growth rate *λ* changed in response to disturbance only by less than 1.5%. This is in part due to the proportional feedback component, which can compensate for the effect of leakiness in biological systems [46]. Indeed, if we assume that *p*_*act*_ synthesis is unaffected by resource couplings, which means that there is no proportional feedback component, the adaptation errors increase (Supplementary Figure S4A-E). Simulations also reveal how changing our circuit’s parameters can improve the controller’s performance. For instance, Figure 5G shows how increasing the amplifier gain *χ* (i.e., the maximum amplifier mRNA production rate) lowers the controller’s adaptation error even as the magnitude of the disturbance rises. Increasing *κ*, the transcription rate of the actuator and the annihilator RNAs has been shown to reduce the adaptation error of the controlled variable’s mean value [44] – albeit in some cases (e.g., the one displayed in Supplementary Figure S4F-J) this can lead to instability [46, 47].

Indirect resource couplings contribute to the breakdown of the assumption of modularity in synthetic biology [12]. However, our controller counters this and renders the expression of different genes more independent of each other, since it keeps ribosome availability in the cell roughly constant. To demonstrate this, we considered how the expression of a heterologous disturbing gene *dist* affects the steady-state concentration of an output protein, encoded by the heterologous gene *x* (Figure 5H). Indeed, the presence of our proportional-integral controller reduced the effect of indirect couplings on the value of *p*_*x*_ almost threefold, as shown in Figure 5I. In the same figure we also mark the estimated boundary of the controller’s operation range, at which the two sides of Equation (31) are strictly equal. Supporting our estimate’s accuracy, this coincides with the point at which increasing the disturbance’s magnitude starts making the output protein’s concentration fall at the same rate both with and without the controller – that is, the point beyond which our controller no longer mitigates burden.

Besides improving the modularity of synthetic biology designs, our controller could slow the loss of synthetic genes by engineered cells. A loss of heterologous gene expression to mutation normally means that the growth rate is less impaired by gene expression burden, which leads to the mutants outcompeting the original engineered cells [48]. Conversely, as long as the controller itself remains functional, it will keep the level of resource competition roughly constant regardless of whether any other synthetic genes are expressed. Without the reduction of translational burden, mutant cells will lack the competitive advantage over the original cells.

## 3 Discussion

The influence of resource competition on synthetic circuit performance can be approached from several perspectives. On one hand, the realistic assumption of fast resource-substrate binding kinetics allows to define the effective rate constants for resource-limited reactions so as to capture the effect of indirect couplings. Simple gene expression models obtained this way can provide easily interpretable predictions of a synthetic circuit’s behavior and the impact of changing its design parameters. However, most such analyses have heretofore been ignoring the wider context of the host organism, neglecting the interplay between heterologous gene expression and the cell’s growth and resource availability [6, 9, 12]. On the other hand, models of the whole bacterial cell allow to simulate how resources are allocated between the host’s own genes and those of the synthetic circuit [25]. Nevertheless, models that involve a large number of variables and parameters are highly complex. This makes the effect of a given design choice difficult to understand without numerically simulating a synthetic circuit’s performance for every set of parameter values considered. Conversely, extensive coarse-graining of cellular processes may abstract the aspects of gene expression that are important for designing gene circuits.

The cell model that we propose aims to combine the strengths of these two approaches. To this end, we apply an established resource-aware modeling framework, based on the use of effective reaction rate constants [9, 12], to a coarse-grained model which relates protein synthesis to cell growth via the finite proteome trade-off [27] and captures near-optimal resource allocation in bacteria by incorporating the principles of flux-parity regulation of gene expression [21]. Spanning only six variables, our host cell model still distinctly considers transcription and translation. For instance, this allows to model RNA-based regulation, which would be impossible with heavily coarse-grained cell models that treat protein production as a single step [20, 21].

The expression of a synthetic circuit by the cell can be easily modeled by introducing additional ODEs mirroring Equations (17)-(18) for each synthetic gene. Particularly, our Matlab implementation of the model, found at https://github.com/KSechkar/rc_e_coli, provides a template .m file, filling which allows to describe an arbitrary synthetic circuit. The resultant script can then be loaded to simulate the circuit’s behavior in the cell, enabling rapid and cheap prototyping of resource-aware biomolecular circuits and controllers *in silico* [49]. Moreover, our modeling framework’s simplicity allows to obtain a range of different analytical relations between a synthetic circuit’s design parameters and the host cell’s growth rate, which have been identified in earlier experimental studies. Consequently, our model provides these relations with a unifying framework. As an example application of our model, we developed and numerically tested a proportional-integral controller that mitigates fluctuations of the host cell’s free ribosome pool size. The circuit senses resource competition not directly but via the changes in a sensor gene’s concentration, meaning that the ribosome availability is inferred via a proxy. However, our model allows to estimate the controller’s set point for the extent of resource competition, as well as the cell growth rate maintained by it and the circuit’s operation range. The fact that the proposed controller acts at a cell-wide level makes our host-aware modeling framework more suited for its analysis than simpler resource competition models that do not consider the cellular context.

The reliability of our model is supported by the concordance of its predictions with a vast body of experimental data from prior studies. Although the parameter values were chosen to capture the experimental characteristics of *E. coli* cells, the finite proteome trade-off is believed to be common for most growing bacterial cells [27], while several other bacteria, such as *S. coelicolor*, have been observed to exhibit behavior consistent with flux-parity regulation of gene expression [21, 50]. This makes our framework potentially applicable to other species upon re-parameterization. The model can also be extended to account for the aspects of cell physiology which are currently omitted but may be particularly relevant for certain applications. For instance, transcriptional resource couplings may become important in the cases of very high heterologous mRNA production. They can be captured by adopting the effective rate constant approach to simplify competitive RNA polymerase binding dynamics similarly to how we considered ribosome pool couplings [12]. Other presently neglected factors, like the heterologous proteins’ metabolic function and toxicity [51], as well as active protein degradation by proteases [52], may be accounted for by introducing additional terms to model equations.

We intend our proposed framework to encourage further exploration of bacterial resource-aware controller designs and to open up avenues for holistic analysis of biocircuit performance in the context of the host cell. A promising research direction is the expansion of bacterial resource-aware control to applications beyond the mitigation of unwanted resource competition interactions, as suggested only recently for mammalian cells in [15]. A potential advantage of resource-aware architectures is a reduced number of direct biological interactions to be engineered, such as gene transcription regulation by transcription factor proteins. Indeed, this has been hinted at by the example of our proportional-integral controller, where the susceptibility of the actuator gene’s transcription rate to resource couplings introduced a proportional feedback component, increasing the controller’s robustness to leakiness without the need for introducing any additional genes.

In conclusion, our work presents a case for combining the insights from resource competition analysis with the appreciation of resource allocation in bacterial cells, which makes it an important step towards easy design of reliable resource coupling-based controllers. Our model’s predictive power can be further enhanced by considering more types of couplings via different resource pools and incorporating the results of future experiments characterising cell physiology. By providing numerical and analytical tools for developing resource-aware circuits with an appreciation for their interactions with the host, we hope to enable the exploration of novel controller designs that leverage indirect couplings.

## 4 Materials and Methods

### 4.1 Numerical Simulations

Our Matlab R2022a model implementation, along with all other scripts used to obtain the results described here, can be found at https://github.com/KSechkar/rc_e_coli. The simulations were run using Matlab’s `ode15s` solver on a laptop with a 6x Intel^®^ Core™ i7-10750H CPU 2.60GHz processor with 16GB RAM, running on Windows 10 21H2.

### 4.2 Parameter Fitting

Parameter fitting was used to estimate the maximum tRNA aminoacylation rate *ν*_*max*_, as well as *K*_*ϵ*_ and *K*_*ν*_, the Michaelis constants determining the translation elongation and tRNA aminoacylation rates. While the metabolic and ribosomal genes’ promoter strengths *α*_*a*_ and *α*_*r*_ also had to be determined this way, we found that the optimality of the fits remained approximately unchanged over a wide range of values as long as the *α*_*r*_:*α*_*a*_ ratio was the same (Supplementary Figure S1B). Therefore, we defined *α*_*a*_ using an order-of-magnitude estimate and fitted this ratio’s value to experimental data. As we outline in Supplementary Information Section 2.3, this then allowed us to obtain more accurate promoter strengths by matching our model’s predictions to experimentally measured RNA production rates [53]. Moreover, since the mutual maximization of tRNA charging and protein synthesis fluxes is known to be achieved when *K*_*ϵ*_ ≈ *K*_*ν*_ [21], we assumed them to be equal, reducing the number of parameter values to be fitted down to only four.

Experimental datapoints for fitting were taken from the study by Scott et al. [18], which measured steady-state growth rates and RNA:protein mass ratios of *E. coli* subjected to different concentrations of the translation-inhibiting drug chloramphenicol in various culture media. In order to convert RNA:protein mass ratios into ribosomal mass fractions, they were multiplied by a conversion factor of 0.4558 [21]. Due to our model’s lack of consideration of the cell’s metabolic regulation mechanisms at very low growth rates, only the datapoints with *λ >* 0.3 were used for parameter fitting.

Similarly to Weisse et al. [27], who fitted their model’s parameters to the same measurements, we estimated different media’s nutrient qualities as six points equally spaced on the logarithmic scale between *σ* = 0.08 and *σ* = 0.5. The effect of chloramphenicol on the cell was modeled by Equations S46-S51, derived and explained in Supplementary Information Section 2.1. Briefly, the ODEs were obtained similarly to the original cell model, first defining a model that explicitly considers all reactions involved in competitive ribosome binding then simplifying the translation dynamics. However, as chloramphenicol binds and disables translating ribosomes, extra terms were introduced to the definitions of apparent mRNA-ribosome dissociation constants {*k*_*i*_}, as well as to the ODE for *R*, redefined as the concentration of the cell’s *operational* ribosomes. As the cell’s measured ribosome content also includes inactivated ribosomes, their levels had to be considered and were denoted as a new variable *B*_*cm*_, whose ODE was added to the model (Supplementary Information Section 2.1).

Fitting was performed using the DiffeRential Evolution Adaptive Metropolis (DREAM) algorithm, which is a variation of the Markov Chain Monte Carlo scheme for inferring parameters’ probability distribution given a set of experimental observations. This involves following the trajectory of a Markov chain, whose states are sets of possible parameter values and whose stationary distribution is equal to the probability distribution of interest. Generally, such a chain is simulated by choosing a potential next set of parameter values according to some proposal distribution and then either accepting or rejecting it based on the probability of observing a known outcome if they are true. In the DREAM modification, the simulation’s efficiency is increased by continuously adapting the proposal distribution based on the states considered in the past, as well as tracking several (in our case, 10) trajectories in parallel [54]. The DREAM simulation was run for 20,000 steps on a server with 2x Intel^®^ Xeon^®^ CPU E5-2670 2.60GHz processors with 128 GB RAM, running on Ubuntu 18.04 LTS; Matlab’s Parallel Computing Toolbox v7.6 was used to track the 10 trajectories simultaneously. The mode of the fitted probability distributions provided the parameter values for our model. The parameters’ starting values and other simulation details can be found in Supplementary Information Section 2.2.

## Data and software availability

All Matlab code and experimental data used in this study described here are available at https://github.com/KSechkar/rc_e_coli.

## Acknowledgements

GBS gratefully acknowledges funding from the Royal Academy of Engineering through the RAE Chair in Emerging Technologies programme (RAEng CiET 1819\5).

## Conflict of interest

The authors declare that there is no conflict of interest.

## Notes

https://github.com/KSechkar/rc_e_coli

### Competing Interest Statement

The authors have declared no competing interest.

