## Supplementary Information for "A coarse-grained bacterial cell model for resource-aware analysis and design of synthetic gene circuits"

### 1 Model Definition

In this section, we derive the simplified model of the host bacterial cell, described by Equations (1)-(6) in the main text's Results section. To this end, we first define a coarse-grained mechanistic cell model that explicitly considers the competitive binding of ribosomes by different mRNAs present in the cell, then reduce the number of considered variables by using the Quasi-Steady-State (QSS) approximation. Finally, in order to further simplify the model, we show how it is possible to omit the Ordinary Differential Equations (ODEs) describing the expression of one of the gene classes into which we partition the bacterium's genome.

#### 1.1 Unsimplified Mechanistic Cell Model with Flux- Parity Regulation

Our model does not take into account transcriptional resource couplings, as they are believed to have a negligible effect on gene expression [1]. Hence, we focus on the competitive binding of a finite pool of ribosomes by mRNAs. We consider three types of native genes in the bacterial cell: metabolic ($a$), ribosomal ($r$), and housekeeping ($q$). For the gene $i \in \{q, a, r\}$, $m_i$ signifies the concentration of its free transcript, $b_i$ the concentration of its mRNA-ribosome complex, and $p_i$ the concentration of the corresponding protein (the number of amino acid residues in one molecule of protein $p_i$ is denoted as $n_i$). However, since ribosomes can be either free or bound by an mRNA, instead of modeling the overall concentration of ribosomal protein $p_r$, we adopt a more specific approach for them, denoting the concentration of *non-translating* ribosomes by $r$. Therefore, the total ribosome concentration can be calculated from the modeled variables as

$$R = r + \sum_{j \in \{q,a,r\}} b_j \tag{S1}$$

Additionally, we look at the pool of protein precursors – aminoacylated (charged) tRNA molecules $t^c$ – as well as uncharged tRNAs $t^u$. The conversion of $t^u$ into $t^c$ is catalysed by the metabolic proteins $p_a$. The set of all reactions that we model is therefore as follows:

- **Transcription:** $\varnothing \xrightarrow{F_i \alpha_i c_i \lambda} m_i$ for $i \in \{q, a, r\}$
- **mRNA degradation and dilution:** $m_i \xrightarrow{\beta_i + \lambda} \varnothing$ for $i \in \{q, a, r\}$

- **mRNA-ribosome binding:** $m_i + r \underset{k_i^+}{\overset{k_i^-}{\rightleftharpoons}} b_i \quad \text{for } i \in \{q, a, r\}$

- **mRNA-ribosome complex dilution:** $b_i \xrightarrow{\lambda} \varnothing \quad \text{for } i \in \{q, a, r\}$

- **Translation:** $b_i + n_i \times t^c \xrightarrow{\epsilon/n_i} m_i + r + p_i + n_i \times t^u \quad \text{for } i \in \{q, a\}$
  but denoted as $b_r + n_r \times t^c \xrightarrow{\epsilon/n_r} m_r + 2 \times r + n_r \times t^u \quad \text{for ribosomal genes}$

- **Protein dilution:** $p_i \xrightarrow{\lambda} \varnothing \quad \text{for } i \in \{q, a\}$
  but denoted as $r \xrightarrow{\lambda} \varnothing \quad \text{for ribosomes}$

- **tRNA synthesis:** $\varnothing \xrightarrow{\psi} t^u$

- **tRNA aminoacylation:** $t^u \xrightarrow{\nu p_a} t^c$

These reactions give rise to the following ODE model for the dynamics of the species' concentrations:

$$\dot{m_i} = F_i c_i \alpha_i \lambda(\epsilon, B) - (\beta_i + \lambda(\epsilon, B)) m_i - k_i^+ m_i r + k_i^- b_i + \frac{\epsilon(t^c)}{n_i} b_i \qquad \text{for } i \in \{q, a, r\} \quad \text{(S2)}$$

$$\dot{b_i} = k_i^+ m_i r - k_i^- b_i - \frac{\epsilon(t^c)}{n_i} b_i - \lambda(\epsilon, B) \cdot b_i \qquad \text{for } i \in \{q, a, r\} \quad \text{(S3)}$$

$$\dot{p_i} = \frac{\epsilon(t^c)}{n_i} b_i - \lambda(\epsilon, B) \cdot p_i \qquad \text{for } i \in \{q, a\} \quad \text{(S4)}$$

$$\dot{r} = \frac{\epsilon(t^c)}{n_r} b_r - \lambda(\epsilon, B) \cdot r + \sum_{j \in \{q,a,r\}} \left( (\frac{\epsilon(t^c)}{n_j} + k_j^-) b_j - k_j^+ m_j r \right) \quad \text{(S5)}$$

$$\dot{t^c} = \nu(t^u, s) \cdot p_a - \epsilon(t^c) \cdot B - \lambda(\epsilon, B) \cdot t^c \quad \text{(S6)}$$

$$\dot{t^u} = \psi(T) - \nu(t^u, \sigma) \cdot p_a + \epsilon(t^c) \cdot B - \lambda(\epsilon, B) \cdot t^u \quad \text{(S7)}$$

(Model I)

Table S1 displays the values, as well as the physiological meaning, of all the parameters used in (Model I) and defining the rates of the reactions listed above. The formulae for the regulation functions and reaction rates, as well as their significance, are given in Table S2. Notably, the expression for $\lambda$ enforces the constant cell mass trade-off identified by Weisse et al. [2]: since the total mass of protein in the cell is by definition

$M = n_r R + \sum_{j\in\{q,a\}} n_j p_j$, the mass of protein in the cell is always constant since:

$$\dot{M} = n_r \dot{R} + \sum_{j\in\{q,a\}} n_j \dot{p_j} \Leftrightarrow \dot{M} = n_r \sum_{j\in\{q,a\}} \dot{b_j} + n_r \dot{r} + \sum_{j\in\{q,a\}} n_j \dot{p_j} \Leftrightarrow$$
$$\Leftrightarrow \dot{M} = \sum_{j\in\{q,a,r\}} n_j \left(\frac{\epsilon}{n_j} b_j\right) - \lambda \left(n_r R + \sum_{j\in\{q,a\}} n_j p_j\right) \Leftrightarrow$$
$$\Leftrightarrow \dot{M} = \epsilon B - \lambda M \Leftrightarrow$$
$$\Leftrightarrow \dot{M} = \epsilon B - \frac{\epsilon B}{M} \cdot M = 0 \tag{S8}$$

Table S1: Parameters appearing in the host cell model ODEs and Table S2's formulae.

| Param. | Description | Value | Units* | Source |
|---|---|---|---|---|
| $M$ | Total cell mass (in amino acids) | $1.19 \cdot 10^9$ | $aa^{\#}$ | [3] |
| $\sigma$ | Extracellular nutrient quality | From 0 to 1 | None | – |
| $\overline{\phi}_q$ | Housekeeping prot. mass fraction§ | 0.59 | None | [4] |
| | **Reaction rates** | | | |
| $\epsilon_{max}$ | Max. translation elongation rate | 72,000 | $aa^{\#}/h$ | [5] |
| $\nu_{max}$ | Max. tRNA charging rate | 4,331.1 | $nM/h$ | Fitted |
| $\psi_{max}$ | Max. tRNA synthesis rate | $1.08 \cdot 10^6$ | $nM/h$ | [6] |
| $K_\epsilon$ | Michaelis-Menten constant for translation elongation rate | 6475.7 | $nM$ | Fitted |
| $K_\nu$ | Michaelis-Menten constant for tRNA charging rate | 6475.7 | $nM$ | Fitted |
| $\tau$ | Michaelis constant for ppGpp signaling (determines $\psi$ and $F_r$) | 1 | None | [6] |
| | **Gene expression** | | | |
| $c_i$ | Concentration of gene $i$ DNA‡ | 1 | $nM$ | Convention |
| $\alpha_q$ | Promoter strength$^{\aleph}$ for gene $q$ | Unspecified§ | None | – |
| $\alpha_a$ | Promoter strength$^{\aleph}$ for gene $a$ | $2.706 \cdot 10^5$ | None | Fitted |
| $\alpha_r$ | Promoter strength$^{\aleph}$ for gene $r$ | $2.189 \cdot 10^5$ | None | Fitted |
| $\beta_i$ | mRNA degradation rate‡ | 6 | $h^{-1}$ | [2] |
| $k_i^+$ | mRNA-ribosome binding rate‡ | 60 | $\frac{1}{nM \cdot h}$ | [2] |
| $k_i^-$ | mRNA-ribosome dissociation rate‡ | 60 | $h^{-1}$ | [2] |
| $n_q$ | Number of amino acids in protein $p_q$ | 300 | $aa^{\#}/nM$ | [2] |
| $n_a$ | Number of amino acids in protein $p_a$ | 300 | $aa^{\#}/nM$ | [2] |
| $n_r$ | Number of amino acids in rib. protein | 7,459 | $aa^{\#}/nM$ | [2] |

**E. coli* volume is $\approx 10^{-18}$ m$^3$, so 1 nM is roughly equivalent to 1 molecule/cell [7].

#Amino acid residues. ‡ Identical across all native genes $i \in \{q, a, r\}$.

§The final simplified (Model III) avoids explicitly modeling housekeeping genes.

$^{\aleph}$ Promoter strength measured as the maximum mRNA synthesis rate per gene copy divided by the cell growth rate at time of measurement.

Table S2: Functions and notations appearing in the host cell model ODEs.

| Func./Not. | Description | Formula | Units* |
|---|---|---|---|
| $B$ | Number of translating ribosomes | $\sum_{j\in\{q,a,r\}} b_j$ | $nM$ |
| $T$ | Proxy for ppGpp concentration (inversely proportional to the alarmone's level) | $t^c/t^u$ | $nM$ |
| **Reaction rate constants** | | | |
| $\epsilon$ | Translation elongation rate | $\epsilon_{max} \cdot \frac{t^c}{t^c+K_\epsilon}$ | $aa^{\#}/h$ |
| $\lambda$ | Growth/dilution rate | $\frac{\epsilon B}{M}$ | $h^{-1}$ |
| $\psi$ | tRNA synthesis rate | $\psi_{max} \cdot \frac{T}{T+\tau}$ | $nM \cdot h^{-1}$ |
| $\nu$ | tRNA charging rate | $\nu_{max} \cdot \sigma \cdot \frac{t^u}{t^u+K_\nu} \cdot p_a$ | $h^{-1}$ |
| **Transcription regulation functions** | | | |
| $F_q$ | Transcription regulation for housekeeping genes | Unspecified§ | None |
| $F_a$ | Transcription regulation for metabolic genes (constitutive) | 1 | None |
| $F_r$ | Transcription regulation for ribosomal genes (via ppGpp) | $\frac{T}{T+\tau}$ | None |

**E. coli* volume is $\approx 10^{-18}$ m$^3$, so 1 nM is roughly equivalent to 1 molecule/cell [7].

#Amino acid residues.

§Defining $F_q$ is not necessary because the final, simplified (Model III) avoids explicitly modeling housekeeping gene expression, as shown in Section 1.5.

#### 1.2 The Quasi-Steady-State Approximation

We now apply the QSS approximation to (Model I) by appreciating that mRNA-ribosome binding and unbinding occur on a much faster timescale than other processes in the cell [7]. Therefore, the mRNA-ribosome complex concentrations can be assumed to be in *quasi-steady state*. Hence, for all $i \in \{q, a, r\}$ we have:

$$\dot{b_i} = 0 \Leftrightarrow k_i^+ m_i r - k_i^- b_i - \frac{\epsilon(t^c)}{n_i} b_i - \lambda(\epsilon, B) \cdot b_i = 0 \Leftrightarrow$$

$$\Leftrightarrow b_i = m_i r \cdot \left( \frac{k_i^- + \epsilon(t^c)/n_i + \lambda(\epsilon, B)}{k_i^+} \right)^{-1} = \frac{m_i}{k_i(\epsilon, B)} \cdot r \tag{S9}$$

#### 1.3 Neglecting the mRNA-Ribosome Complex Dilution

The retrieval of the mRNA-ribosome complexes' concentrations using Equation (S9) is nevertheless hindered by the dilution term $\lambda(\epsilon, B)$ in $k_i(\epsilon, B)$, which itself depends on the total concentration of translating ribosomes. Instead of the full expression for $k_i(\epsilon, B)$, it would therefore be desirable to use an approximation given by Equation (S10), which neglects dilution due to cell growth and thus always slightly underestimates $k_i$.

$$k_i(\epsilon, B) \approx k_i(\epsilon) = \frac{k_i^- + \epsilon/n_i}{k_i^+} \tag{S10}$$

Let us now show that in all realistic scenarios this approximation is reasonably close to the actual value of $k_i(\epsilon, B)$. First, we substitute the formula for $\lambda$ from Table S2 into the full expression for $k_i(\epsilon, B)$ from Equation (S9), obtaining:

$$k_i(\epsilon, B) = \frac{k_i^- + \frac{\epsilon(t^c)}{n_i} + \lambda(\epsilon, B)}{k_i^+} = \frac{k_i^- + \frac{\epsilon(t^c)}{n_i} + \frac{\epsilon B}{M}}{k_i^+} = \frac{k_i^-}{k_i^+} + \frac{\epsilon}{k_i^+} \left( \frac{1}{n_i} + \frac{B}{M} \right) \tag{S11}$$

The effect of growth on this value is thus restricted to the $^B/_M$ component of the $\epsilon$-dependent term of the expression. Now, let us consider the mass fraction of all ribosomal proteins in the cell $\phi_r = \frac{n_r R}{M}$. In the extensive experimental studies of E. coli growth conducted up to this point, it has not been found to exceed 0.3 [6]. Therefore, noticing that $B \leq R$ by definition, for all realistic purposes we can impose a cap on $^B/_M$:

$$\frac{B}{M} \leq \frac{R}{M} = \frac{\phi_r}{n_r} \leq \frac{0.3}{n_r} \tag{S12}$$

The metabolic and housekeeping proteins are assumed to span $n_q = n_a = 300$ amino acids, while a single ribosome comprises $n_r = 7549$ residues (see Table S1). For the former two species we thus have

$$\frac{1}{n_q} = \frac{1}{n_a} = \frac{1}{300} \approx 3.33 \times 10^{-3} \gg 4.02 \times 10^{-5} = \frac{0.3}{7459} = \frac{0.3}{n_r} \geq \frac{B}{M} \tag{S13}$$

which means that the contribution of dilution to the $\epsilon$-dependent term can be safely disregarded, vindicating the use of an approximation from Equation (S10).

Nonetheless, the situation is more complicated in the case of ribosomes, as for $n_i = n_r$ the $B/M$ component can reach up to 30% of $1/n_r$. However, for such a high number of amino acid residues, the whole $\epsilon$-dependent term contributes very little to $k_r$, even at the highest translation rates possible. This can be demonstrated by calculating the approximation error $\delta_r(\epsilon)$ – that is, the difference between the estimate $k_r(\epsilon)$ and the real value $k_r(\epsilon, B)$, divided by the real value. Since the approximate value of $k_r$ is always lower than the real one, dividing the difference by $k_r(\epsilon)$ instead of $k_r(\epsilon, B)$ bounds the approximation error from above:

$$\delta_r(\epsilon) = \frac{k_r(\epsilon, B) - k_r(\epsilon)}{k_r(\epsilon, B)} \leq \frac{k_r(\epsilon, B) - k_r(\epsilon)}{k_r(\epsilon)} = \frac{\frac{\epsilon}{k_r^+} \cdot \frac{0.3}{n_r}}{\frac{k_r^-}{k_r^+} + \frac{\epsilon}{k_r^+ n_r}} = \frac{0.3\epsilon}{k_r^- n_r + \epsilon} = \delta_r^{bnd}(\epsilon) \tag{S14}$$

It is easy to see that for positive translation elongation rates this bound increases monotonously with $\epsilon$, so to cap the error for all cases it is enough to evaluate $\delta_r^{bnd}(\epsilon)$ for the maximum possible translation elongation rate. Using the value from Table S1, we find this cap to be merely

$$\delta_r^{bnd}(\epsilon_{max}) = \frac{0.3\epsilon_{max}}{k_r^- n_r + \epsilon_{max}} = 4.15\% \tag{S15}$$

In summary, we have shown that for any transcript, it is possible to reliably approximate its lumped "mRNA-ribosome dissociation constant" $k_i(\epsilon, B)$ by an expression $k_i(\epsilon)$ that does not acknowledge the effects of dilution caused by cell growth, removing this value's dependence on the concentration of translating ribosomes (Equation (S11)). As shown in the next section, this allows to greatly simplify the analysis of competitive ribosome binding by different mRNAs. From this point on, we will be denoting the approximate value $k_i(\epsilon)$ simply as $k_i$ to avoid clutter. The number of ribosomes translating gene $i$'s mRNA can therefore be denoted as

$$b_i \approx \frac{m_i}{k_i} r \tag{S16}$$

#### 1.4 Cell Model with the QSS Approximation

Let us now remember the definition for the total number of ribosomes in the cell $R$ from Equation (S1). Together with Equation (S16), it yields

$$R = \sum_{\forall i} b_i + r = r(1 + \sum_{\forall i} \frac{m_i}{k_i}) \Leftrightarrow r = R/(1 + \sum_{\forall i} \frac{m_i}{k_i}) \Rightarrow$$

$$\Rightarrow b_i = \frac{m_i/k_i}{1 + \sum_{j \in \{q,a,r\}} m_j/k_j} R = \frac{m_i/k_i}{D} R \tag{S17}$$

where

$$D = 1 + \sum_{j \in \{q,a,r\}} m_j/k_j \tag{S18}$$

Consequently, we can also state that

$$B = \sum_{\forall i} b_i = \frac{\sum_{j \in \{q,a,r\}} m_j/k_j}{1 + \sum_{j \in \{q,a,r\}} m_j/k_j} R = (1 - \frac{1}{D}) R \tag{S19}$$

By providing the expressions for $\{b_i\}$ and $B$, Equations (S17) and (S19) rid us of the necessity to explicitly consider mRNA-ribosome binding and treat free and translating ribosomes as separate species. Moreover, with the complex dilution considered insignificant, the sum of Equation (S2)'s last three terms is roughly equal to $-\dot{b_i}$, which is zero due to the QSS assumption. This yields us a simplified (Model II). Importantly, Equation (S22) now gives an ODE for the total number of ribosomes $R$ and not just the free ribosome count $r$.

$$\dot{m_i} = F_i c_i \alpha_i \lambda(\epsilon, B) - (\beta_i + \lambda(\epsilon, B)) m_i \quad \text{for } i \in \{q,a,r\} \tag{S20}$$

$$\dot{p_i} = \frac{\epsilon(t^c)}{n_i} \cdot \frac{m_i/k_i}{1 + \sum_{j \in \{q,a,r\}} m_j/k_j} R - \lambda(\epsilon, B) \cdot p_i \quad \text{for } i \in \{q,a,r\} \tag{S21}$$

$$\dot{R} = \frac{\epsilon(t^c)}{n_r} b_r - \lambda(\epsilon, B) \cdot r \tag{S22}$$

$$\dot{t^c} = \nu(t^u, s) \cdot p_a - \epsilon(t^c) \cdot B - \lambda(\epsilon, B) \cdot t^c \tag{S23}$$

$$\dot{t^u} = \psi(T) - \nu(t^u, \sigma) \cdot p_a + \epsilon(t^c) \cdot B - \lambda(\epsilon, B) \cdot t^u \tag{S24}$$

(Model II)

If the system is in steady state (which here is indicated by drawing bars over the variables), a useful expression for a given gene's equilibrium protein concentration $\overline{p}_i$ can be derived as shown below.

$$\dot{p}_i = 0 \Leftrightarrow \frac{\overline{\epsilon}}{n_i} \cdot \frac{\overline{m}_i/\overline{k}_i}{\overline{D}}\overline{R} - \overline{\lambda}\overline{p}_i = 0 \Leftrightarrow$$

$$\Leftrightarrow \overline{p}_i = \frac{M}{n_i} \cdot \frac{\overline{m}_i/\overline{k}_i}{\sum_{j\in\{q,a,r\}} \overline{m}_j/\overline{k}_j} \tag{S25}$$

Importantly, Equation (S25) is merely a *relation* between the steady-state mRNA and protein concentrations, and does not allow to analytically determine $\overline{p}_i$ straightaway. This is because the formulae for $k_i$ values include the translation elongation rate $\epsilon$, while the steady-state mRNA concentrations $\overline{m}_i$ depend on the cell growth rate $\lambda$. Since $\epsilon$ and $\lambda$ are not constant and depend on the cell's state, the values of $\overline{m}_i$ and $\overline{k}_i$ cannot be found based on Equation (S25) alone.

Let us conclude this section by stating a useful relation for $\overline{\phi}_i$, the steady-state mass fraction of protein $p_i$. Multiplying the protein's concentration by one molecule's weight in amino acid residues and dividing it by the cell's total protein mass, we get:

$$\overline{\phi}_i = \frac{n_i\overline{p}_i}{M} = \frac{\overline{m}_i/\overline{k}_i}{\sum_{j\in\{q,a,r\}} \overline{m}_j/\overline{k}_j} \tag{S26}$$

#### 1.5 Neglecting the Housekeeping Genes

The key property of housekeeping genes is that their expression remains constant and growth rate-independent. In fact, the assumption that $\phi_q$ is unchanging regardless of the growth rate or nutrient availability has been shown to hold under a wide range of conditions [4], yielding reliable predictions about the dynamics of cellular processes and not just their steady states [6]. Therefore, instead of explicitly modeling transcription and translation of the housekeeping genes, we can further simplify the model by assuming that $\phi_q \equiv \overline{\phi}_q \equiv 0.59$ (and thus $p_q \equiv \overline{p}_q \equiv \frac{\overline{\phi}_q M}{n_q}$) in all cases [4]. Therefore, at any point in time

$$\frac{m_q/k_q}{\sum_{j\in\{q,a,r\}} m_j/k_j} = \overline{\phi}_q \tag{S27}$$

This in turn means that

$$\sum_{j\in\{q,a,r\}} m_j/k_j = \frac{1}{1-\overline{\phi}_q} \sum_{j\in\{a,r\}} m_j/k_j \tag{S28}$$

and

$$D = 1 + \frac{1}{1-\overline{\phi}_q} \sum_{j\in\{a,r\}} m_j/k_j \tag{S29}$$

$B$, the total concentration of mRNA-ribosome complexes, including those with the houskeeping gene mRNA, can still be found from $D$ as described by Equation (S19). Consequently, it is possible to model ribosomal and metabolic protein expression, cell growth, and tRNA charging and synthesis without knowing the exact mRNA concentrations for the housekeeping genes. Hence, we can modify (Model II) to obtain (Model III). This is the simplified mechanistic cell modeling framework that we describe in the article's main text (Equations (1)-(6) in Results) and employ to obtain the results presented in this paper.

$$\dot{m_i} = F_i c_i \alpha_i \lambda(\epsilon, B) - (\beta_i + \lambda(\epsilon, B)) m_i \qquad \text{for } i \in \{a, r\} \tag{S30}$$

$$\dot{p_a} = \frac{\epsilon(t^c)}{n_a} \cdot \frac{m_a/k_a}{1 + \frac{1}{1-\overline{\phi}_q} \sum_{j\in\{a,r\}} m_j/k_j} R - \lambda(\epsilon, B) \cdot p_a \tag{S31}$$

$$\dot{R} = \frac{\epsilon(t^c)}{n_r} \cdot \frac{m_r/k_r}{1 + \frac{1}{1-\overline{\phi}_q} \sum_{j\in\{a,r\}} m_j/k_j} R - \lambda(\epsilon, B) \cdot r \tag{S32}$$

$$\dot{t^c} = \nu(t^u, s) \cdot p_a - \epsilon(t^c) \cdot B - \lambda(\epsilon, B) \cdot t^c \tag{S33}$$

$$\dot{t^u} = \psi(T) - \nu(t^u, \sigma) \cdot p_a + \epsilon(t^c) \cdot B - \lambda(\epsilon, B) \cdot t^u \tag{S34}$$

(Model III)

Using Equation (S28), it is also possible to rewrite Equations (S25) and (S26) to obtain expressions for the steady-state protein concentrations and mass fractions:

$$\overline{p}_i = \frac{M}{n_i} \cdot \frac{\overline{m}_i/\overline{k}_i}{\frac{1}{1-\overline{\phi}_q} \sum_{j\in\{a,r\}} \overline{m}_j/\overline{k}_j} \tag{S35}$$

$$\overline{\phi}_i = \frac{\overline{m}_i/\overline{k}_i}{\frac{1}{1-\overline{\phi}_q} \sum_{j\in\{a,r\}} \overline{m}_j/\overline{k}_j} \tag{S36}$$

### 2 Parameter Fitting

Here, we provide the details of our fitting procedure for the model parameters that were not taken from literature. First, we outline how the cell model was modified to incorporate the effects of the ribosome-inactivating antibiotic chloramphenicol, since it had been present in the culture media of some *E. coli* cells in the experiment whose results we used for fitting [8]. Next, we describe the parameters and outcome of the Markov Chain Monte Carlo (MCMC) simulation that we used to infer the parameter values from data. Finally, we show how the inferred values were postprocessed to make the model more realistic.

#### 2.1 modeling Ribosome Inactivation

In order to determine the parameters' values, we fit our model to experimental data obtained by Scott et al. [8]. In their experiment, the growth rate and the ratio of total RNA mass to the overall mass of protein in the cell were measured for *E. coli* grown in different conditions (for our fitting, the RNA:protein ratios that they recorded were multiplied by a conversion factor of 0.4558 to obtain ribosomal mass fractions [6]). These sundry growth conditions were modeled by varying the culture medium's nutritional quality and the amount of chloramphenicol present in it. Chloramphenicol is an antibiotic that inactivates translating ribosomes by binding them instead of the tRNA that carries the amino acid supposed to be added to the peptide chain. We assume that the presence of chloramphenicol introduces two additional kinds of reactions besides those displayed in Section 1.1:

- **Ribosome inactivation:** $b_i + h \xrightarrow{k_{cm}} B_{cm} \quad \text{for } i \in \{q, a, r\}$
- **Inactivated ribosome dilution:** $B_{cm} \xrightarrow{\lambda} \varnothing$

Here, $b_i$ denotes the mRNA-ribosome complex for gene $i$, $h$ signifies chloramphenicol, and $B_{cm}$ the inactivated ribosome. The first reaction equation stands for ribosome inactivation by chloramphenicol, whereas the second describes the dilution of the inactivated ribosomes due to cell division.

Notably, we do not explicitly model the antibiotic's transport across the cell membrane, so $h$ denotes the concentration of chloramphenicol *in the medium.* The value of the binding rate constant $k_{cm}$ therefore implicitly accounts for conversion between the extracellular and the intracellular antibiotic levels. Moreover, the share of chloramphenicol molecules bound by ribosomes is assumed to be very small compared to the antibiotic's abundance in the media, so we consider $h$ constant.

Then, let $R = r + \sum_{j\in\{q,a,r\}} b_i$ denote exclusively non-inactivated ribosomes, and let $B_{cm}$ signify the concentration of ribosomes bound by chloramphenicol. The experimentally measured ribosome fraction, nevertheless, still includes both operational and disabled ribosomes. Likewise to Section 1.1, we obtain the unsimplified ODE (Model IV).

$$\dot{m_i} = F_i c_i \alpha_i \lambda(\epsilon, B) - (\beta_i + \lambda(\epsilon, B))m_i - k_i^+ m_i r + k_i^- b_i + \frac{\epsilon(t^c)}{n_i} b_i \qquad \text{for } i \in \{q,a,r\} \quad \text{(S37)}$$

$$\dot{b_i} = k_i^+ m_i r - k_i^- b_i - \frac{\epsilon(t^c)}{n_i} b_i - \lambda(\epsilon, B)\cdot b_i - k_{cm} h b_i \qquad \text{for } i \in \{q,a,r\} \quad \text{(S38)}$$

$$\dot{p_i} = \frac{\epsilon(t^c)}{n_i} b_i - \lambda(\epsilon, B)\cdot p_i \qquad \text{for } i \in \{q,a\} \quad \text{(S39)}$$

$$\dot{r} = \frac{\epsilon(t^c)}{n_r} b_r - \lambda(\epsilon, B)\cdot r + \sum_{j\in\{q,a,r\}} \left( (\frac{\epsilon(t^c)}{n_j} + k_j^-) b_j - k_j^+ m_j r \right) \quad \text{(S40)}$$

$$\dot{t^c} = \nu(t^u, s)\cdot p_a - \epsilon(t^c)\cdot B - \lambda(\epsilon, B)\cdot t^c \quad \text{(S41)}$$

$$\dot{t^u} = \psi(T) - \nu(t^u, \sigma)\cdot p_a + \epsilon(t^c)\cdot B - \lambda(\epsilon, B)\cdot t^u \quad \text{(S42)}$$

$$\dot{B}_{cm} = k_{cm} h B - B_{cm}\lambda(\epsilon, B) \quad \text{(S43)}$$

(Model IV)

Applying the QSS approximation to Equation (S38), we obtain

$$b_i = m_i r \cdot \left( \frac{k_i^- + \epsilon(t^c)/n_i + \lambda(\epsilon, B) + k_{cm} h}{k_i^+} \right)^{-1} = \frac{m_i}{k_i(\epsilon, B, h)} r \quad \text{(S44)}$$

An additional non-negative term – that is, $k_{cm}h$ – can only render the growth-dependent component's contribution even less significant relative to other terms. Hence, the considerations outlined in Section 1.3 remain valid and allow us to employ a growth-independent approximation for the mRNA-ribosome dissociation constant. We thus define $\tilde{k_i}$, the approximate mRNA-ribosome dissociation constant with adjustment for chloramphenicol, as

$$\tilde{k_i}(\epsilon, h) = \frac{\epsilon/n_i + k_i^- + k_{cm} h}{k_i^+} \quad \text{(S45)}$$

Consequently, we can simplify (Model IV) analogously to how we reduced (Model I) in Sections 1.4-1.5, this time with $\tilde{k_i}$ instead of $k_i$. This yields us (Model V), which we used in our fitting procedure to predict a cell's behavior for given culturing conditions and parameter values.

$$\dot{m}_i = F_i c_i \alpha_i \lambda(\epsilon, B) - (\beta_i + \lambda(\epsilon, B)) m_i - k_{cm} h \cdot \frac{m_i/\tilde{k}_i}{1 + \frac{1}{1-\bar{\phi}_q} \sum_{j \in \{a,r\}} m_j/\tilde{k}_j} R \quad \text{for } i \in \{a, r\} \tag{S46}$$

$$\dot{p}_a = \frac{\epsilon(t^c)}{n_a} \cdot \frac{m_a/\tilde{k}_a}{1 + \frac{1}{1-\bar{\phi}_q} \sum_{j \in \{a,r\}} m_j/\tilde{k}_j} R - \lambda(\epsilon, B) \cdot p_a \tag{S47}$$

$$\dot{R} = \frac{\epsilon(t^c)}{n_r} \cdot \frac{m_r/\tilde{k}_r}{1 + \frac{1}{1-\bar{\phi}_q} \sum_{j \in \{a,r\}} m_j/\tilde{k}_j} R - \lambda(\epsilon, B) \cdot r - k_{cm} h B \tag{S48}$$

$$\dot{t}^c = \nu(t^u, s) \cdot p_a - \epsilon(t^c) \cdot B - \lambda(\epsilon, B) \cdot t^c \tag{S49}$$

$$\dot{t}^u = \psi(T) - \nu(t^u, \sigma) \cdot p_a + \epsilon(t^c) \cdot B - \lambda(\epsilon, B) \cdot t^u \tag{S50}$$

$$\dot{B}_{cm} = k_{cm} h B - \lambda(\epsilon, B) \cdot B_{cm} \tag{S51}$$

(Model V)

#### 2.2 MCMC fitting

Fitting was required to determine the values of:

- $\nu_{max}$, the maximum tRNA charging rate
- $\alpha_a$ and $\alpha_r$, the metabolic and ribosomal genes' promoter strengths
- $K_\epsilon$ and $K_\nu$, the Michaelis constants determining the translation elongation and tRNA charging rates, respectively

We performed the fitting with the help of the Differential Evolution Adaptive Metropolis (DREAM) algorithm [9], running 10 chains in parallel for 20,000 steps. As a Markov Chain Monte Carlo (MCMC) method, it constructs Markov chains whose states are the possible sets of parameter values and whose stationary distribution equals the probability distribution of these parameter sets. This is achieved by selecting a set of parameter values according to some proposal distribution and accepting it as the next state in the chain with a probability that is proportional to the likelihood of the experimental measurements for this parameter set. In our case, the experimentally measured values of the steady-state growth rate $\overline{\lambda}$ and ribosomal mass fraction $\overline{\phi}_r$ were assumed to have independent normal distributions around the mean values predicted by the model with a given parameter

Table S3: Parameters fitted to experimental data and the possible ranges for their values input to the DREAM algorithm.

| Parameter(s) | Description | Min. value | Max. value | Units |
|---|---|---|---|---|
| $K_\epsilon = K_\nu$ | Michaelis constants determining the translation elongation and tRNA charging rates | 800 | $8 \cdot 10^6$ | $h^{-1}$ |
| $k_{cm}$ | Chloramphenicol binding rate constant | $3.594 \cdot 10^{-6}$ | $3.594 \cdot 10^{-2}$ | $\frac{1}{nM \cdot h}$ |
| $\alpha_r : \alpha_a$ | Ribosomal to metabolic gene promoter strength ratio | 0.01 | 100 | None |
| $\nu_{max}$ | Maximum tRNA charging rate | 60 | $6 \cdot 10^5$ | $nM/h$ |

set. We used the standard deviation values of 0.0447 $h^{-1}$ and 0.018976 for $\overline{\lambda}$ and $\overline{\phi}_r$ respectively, calculated as the average measurement errors across all conditions observed by Scott et al. [8]. In line with [6], the conversion of the RNA:protein mass ratio into ribosomal mass fraction was achieved by multiplying the measurement by 0.4558.

Notably, the model was fitted solely to the datapoints with growth rates above 0.3 $h^{-1}$, because at slower growth rates experimental metabolic regulation is predominated by mechanisms unconsidered by our model, while experimental measurements can be unreliable (see the main text's Results section and [6]). Furthermore, to improve the fitting procedure's efficiency, we reduced the number of values to be determined as follows. First, near-optimal resource allocation is known to be achieved when $K_\epsilon$ and $K_\nu$ are roughly equal [6]. Consequently, we treated both Michaelis constants as a single parameter whose value was to be determined. Second, in our initial MCMC runs, we observed that the likelihood changes very little when the absolute values of $\alpha_a$ and $\alpha_r$ are varied but their ratio stays the same (Figure S1B) . Therefore, instead of fitting both values, we fixed $\alpha$ at a crude order-of-magnitude estimate of $\alpha_a = 2.72 \cdot 10^5$, making the metabolic genes account for a third of all mRNA synthesis in the cell at a reference growth rate of 0.7 $h^{-1}$[3], and used MCMC to determine the $\alpha_r$:$\alpha_a$ ratio. The admissible ranges for the parameter values that we defined for the fitting are displayed in Table S3.

After running the MCMC algorithm, the estimated parameter values were obtained by finding the mode of the inferred posterior probability distribution. We then used these values to improve upon our estimate for $\alpha_a$ as described in Section 2.3.

In order to asses the fit's sloppiness, we estimated the posterior distribution's Fisher Information Matrix (FIM) by retrieving the pseudo-inverse of the variance-covariance matrix [2, 10]. The magnitudes of the FIM's eigenvalues were spread over almost 24 orders of magnitude (Figure S1C), hinting that the model is truly sloppy, i.e., that its behavior is largely independent from the exact values of most parameters [11]. Indeed, as shown in Figure S1D, the estimates of parameter sensitivities based on the FIM [10] reveal that the model is relatively insensitive to changes in most parameter values. An exception to this is the ratio between the metabolic and ribosomal genes' promoter strengths $\alpha_r : \alpha_a$, the sensitivity to which is overwhelmingly large when compared to all other parameters. This can likely be explained by the fact that under our model the cell achieves near-optimal growth rates by managing resource allocation between ribosome and metabolic gene synthesis. Hence, a very different regulatory response may be required from the cell if the transcription rates of the two genes' mRNAs are altered, so the cell's behavior in given conditions may significantly change.

#### 2.3 Scaling the Promoter Strengths

During parameter fitting, we used a crude order-of-magnitude estimate of $\alpha_a = 3.462 \cdot 10^5$. However, with the ratio between $\alpha_a$ and $\alpha_r$ known, the gene transcription rates can be determined with greater accuracy. To this end, we consider the *E. coli* cell in steady state growing at a rate of $\overline{\lambda} \approx 0.7/h$. The total rate of mRNA production in these conditions was experimentally observed to be $A = 1.02 \cdot 10^6 \ nucleotides/min = 6.12 \cdot 10^7 nucleotides/h$ [3]. In the following section, we express this quantity in terms of $\alpha_a$, hence becoming able to deduce the metabolic gene transcription rate.

According to the MCMC fit, the maximum ribosomal gene transcription rate $\alpha_r$ constitutes 0.953427 of $\alpha_a$. Therefore, knowing the value of the ribosomal gene transcription regulation function $\overline{F}_r$ would enable us to find the total ribosomal mRNA production rate as

$$\text{Total } m_r \text{ synthesis rate} = \overline{F}_r \cdot \overline{\lambda} \cdot \alpha_r = \overline{F}_r \cdot \overline{\lambda} \cdot 0.95342 \cdot \alpha_a \quad \text{(S52)}$$

As for the housekeeping genes, we assume the housekeeping and metabolic gene transcripts to have 1) similar degradation rates $\beta_q = \beta_a = 6 \ h^{-1}$ and 2) similar mRNA-ribosome dissociation constants $k_q = k_a$ (due to having the same lengths $n_a = n_q = 300$). Then, by Equation (S26) we have

$$\frac{\overline{\phi}_q}{\overline{\phi}_a} = \frac{\overline{m}_q/\overline{k}_q}{\overline{m}_a/\overline{k}_a} = \frac{\left(\frac{\overline{F}_q c_q \alpha_q \overline{\lambda}}{\beta_q + \overline{\lambda}} \cdot \frac{1}{\overline{k}_q}\right)}{\left(\frac{c_a \alpha_a \overline{\lambda}}{\beta_a + \overline{\lambda}} \cdot \frac{1}{\overline{k}_a}\right)} = \frac{\overline{F}_q c_q \alpha_q}{c_a \alpha_a} \Leftrightarrow$$

$$\Leftrightarrow \text{Total } m_q \text{ synthesis rate} = \frac{\overline{\phi}_q}{\overline{\phi}_a} \cdot \overline{\lambda} \cdot \alpha_a = \frac{0.59}{\overline{\phi}_a} \cdot \overline{\lambda} \cdot \alpha_a \tag{S53}$$

Importantly, our model assumes that a transcript can be bound by one ribosome at a time. In reality, however, multiple ribosomes can translate the same mRNA molecule simultaneously. Likewise to the resource competition model proposed by Qian et al. [7], we implicitly address this discrepancy by the adjusting the mRNA synthesis rates. Namely, the minimum distance between translating ribosomes is estimated at $\approx 25$ codons [12], so a single mRNA molecule that spans $n_i$ codons can be bound by up to $n_i/25$ ribosomes. Hence, the apparent production rate of the mRNA is the actual tramscription rate times $n_i/25$. In combination with our previous considerations, the *real* total mRNA production rate in the cell is the sum of

$$\begin{aligned}
\text{Actual total } m_q \text{ synthesis rate} &= \left(\frac{n_q}{25}\right)^{-1} \cdot \frac{0.59}{\overline{\phi}_a} \cdot \overline{\lambda} \cdot \alpha_a \\
\text{Actual total } m_r \text{ synthesis rate} &= \left(\frac{n_r}{25}\right)^{-1} \cdot \overline{F}_r \cdot 0.953427 \cdot \overline{\lambda} \cdot \alpha_a \\
\text{Actual total } m_a \text{ synthesis rate} &= \left(\frac{n_a}{25}\right)^{-1} \cdot \overline{\lambda} \cdot \alpha_a
\end{aligned}$$

Finally, considering that A is given in nucleotides per minute, the rate of production of a given mRNA $m_i$ should also be multiplied by the number of nucleotides it comprises, i.e., $3n_i$. In summary, all these steps give rise to the following relationship:

$$A = \alpha_a \cdot 0.7 \cdot \left(75 \cdot \frac{0.59}{\overline{\phi}_a} + 75 \cdot \overline{F}_r \cdot 0.953427 + 75\right) \tag{S54}$$

Since model predictions change little upon the scaling of $\alpha_a$ and $\alpha_r$, we start by simulating the model for the fitted parameter values given in Table S3, which allows us to numerically estimate $\overline{F}_r$ and $\overline{\phi}_a$. Using $\sigma = 0.1705$ as a nutrient quality coefficient that yields $\lambda = 0.7004 \approx 0.7\ h^{-1}$, we obtain

$$\overline{\phi}_a \approx 0.309 \text{ and } \overline{F}_r \approx 0.0992$$

And therefore the metabolic protein transcription rate is

$$\alpha_a = 3.881 \cdot 10^5$$

as specified in Table S1. The maximum ribosomal gene transcription rate is therefore obtained by multiplying this value by 0.95342, hence

$$\alpha_r = 3.687 \cdot 10^5$$

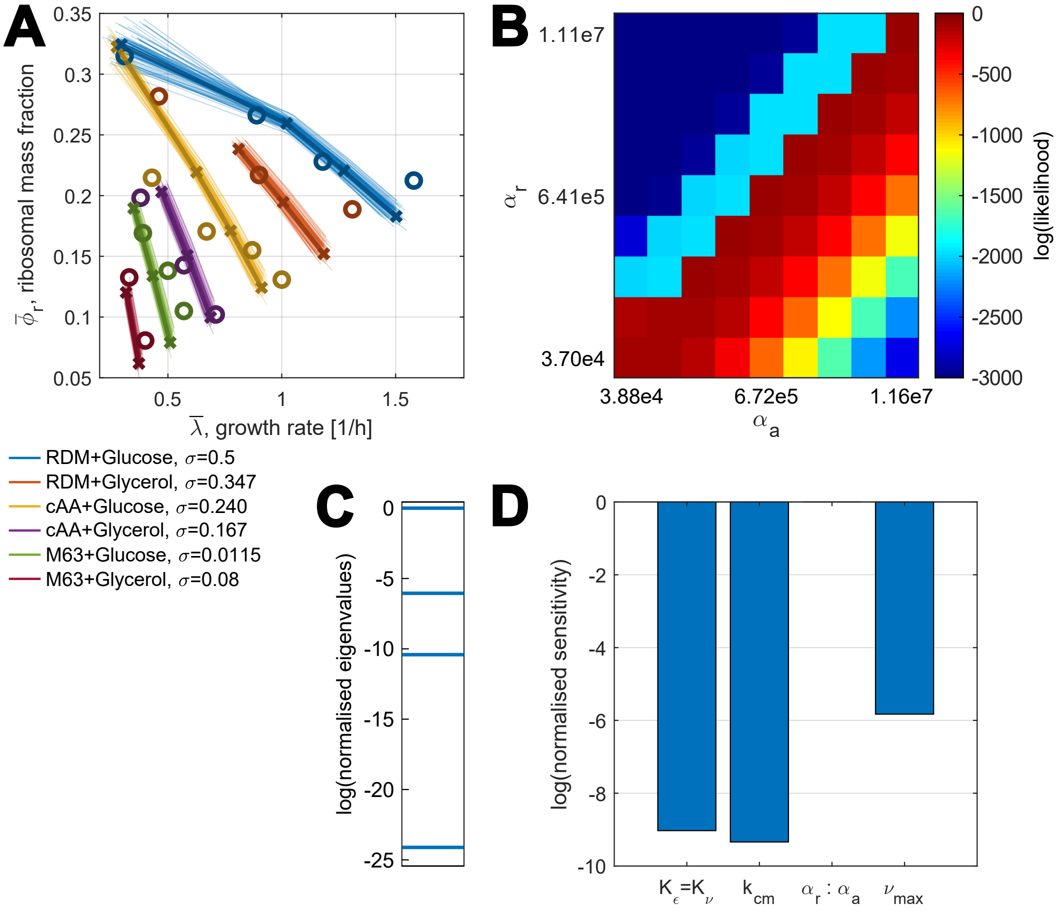

Figure S1: MCMC fitting outcome and parameter sensitivity analysis. **(A)** MCMC fitting outcome for the model. Thin lines: predictions for 100 parameter sets randomly drawn from the obtained probability distribution. Bold dark lines and crosses: model predictions for the maximum a posteriori estimate (found as the distribution's mode) of the parameter values, which we ultimately employ in our model and display in Table S1. Bold circles: experimental measurements by Scott et al. [8] that were used in our fitting procedure. **(B)** Logarithms of experimental measurements' likelihood for different values of metabolic and ribosomal genes' promoter strengths $\alpha_a$ and $\alpha_r$. Values of all other model parameters were taken from Table S1. Observe how consistent the likelihoods stay along the diagonals going through the points that have the same $\alpha_r : \alpha_r$ ratios. **(C)** Logarithms of the FIM's eigenvalues normalized by the largest eigienvalue's magnitude. **(D)** Parameter sensitivities evaluated using the FIM, normalized to yield 1 when summed. The closer is the logarithm of parameter sensitivity to 0, the greater is the effect of the corresponding parameter's value on the model fit.

### 3 Heterologous Gene Expression modeling

In this section, we describe how to extend the host cell model to include the expression of heterologous genes. Then, we derive the analytical relations between heterologous gene expression and the cell's state that we describe in the main text.

#### 3.1 Extending the Mechanistic Cell Model

The simplified mechanistic (Model III), described in Section 1.5, can easily be extended to model the expression of heterologous genes in the bacterium. In addition to the set of the modeled native genes $\{a, r\}$, let there also be a set of heterologous genes $X = \{x_1, x_2 \ldots, x_L\}$. These genes are characterized by the same types of gene expression parameters as the native ones ($\{c_{x_l}\}, \{\alpha_{x_l}\}$, etc.) and each have a corresponding gene- and circuit-specific transcription regulation function $\{F_{x_l}\}$.

The existence of additional genes besides the native ones does not fundamentally change the considerations that enabled the reduction of the full mechanistic (Model I) to the simplified (Model III) outlined in Sections 1.2-1.5, so we can still apply the QSS approximation and avoid explicitly modeling housekeeping gene expression (the fixed housekeeping protein mass fraction can be still assumed to equal $\overline{\phi}_q = 0.59$ [8, 13, 4]). As a result, we use the same general form for the ODEs describing a gene's mRNA concentration $m_i$ and protein concentration $p_i$ as Equations (S30) and (S31) in (Model III). This lets us describe the host cell and the synthetic circuit by (Model VI).

$$\dot{m_i} = F_i c_i \alpha_i \lambda(\epsilon, B) - (\beta_i + \lambda(\epsilon, B)) m_i \qquad \text{for } i \in \{a, r\} \tag{S55}$$

$$\dot{p_a} = \frac{\epsilon(t^c)}{n_a} \cdot \frac{m_a/k_a}{1 + \frac{1}{1-\overline{\phi}_q} \sum_{j \in \{a,r\} \cup X} m_j/k_j} R - \lambda(\epsilon, B) \cdot p_a \tag{S56}$$

$$\dot{R} = \frac{\epsilon(t^c)}{n_r} \cdot \frac{m_r/k_r}{1 + \frac{1}{1-\overline{\phi}_q} \sum_{j \in \{a,r\} \cup X} m_j/k_j} R - \lambda(\epsilon, B) \cdot R \tag{S57}$$

$$\dot{t^c} = \nu(t^u, s) \cdot p_a - \epsilon(t^c) \cdot B - \lambda(\epsilon, B) \cdot t^c \tag{S58}$$

$$\dot{t^u} = \psi(T) - \nu(t^u, \sigma) \cdot p_a + \epsilon(t^c) \cdot B - \lambda(\epsilon, B) \cdot t^u \tag{S59}$$

$$\dot{m_{x_l}} = F_{x_l} \cdot c_{x_l} \alpha_{x_l} - (\beta_{x_l} + \lambda(\epsilon, B)) m_{x_l} \qquad \text{for } x_l \in X \tag{S60}$$

$$\dot{p_{x_l}} = \frac{\epsilon(t^c)}{n_{x_l}} \cdot \frac{m_{x_l}/k_{x_l}}{1 + \frac{1}{1-\overline{\phi}_q} \sum_{j \in \{a,r\} \cup X} m_j/k_j} R - \lambda(\epsilon, B) \cdot p_{x_l} \qquad \text{for } x_l \in X \tag{S61}$$

(Model VI)

(Model VI) allows to numerically simulate the behavior of a cell expressing heterologous proteins, provided that the gene expression burden does not decrease the growth rate enough to activate the cell's stress response mechanisms, which our model does not consider (in the main text's Results section, we postulate a threshold of $\lambda = 0.3\ h^{-1}$, beyond which our model's predictions significantly diverge with experimental measurements). Moreover, by solving $\dot{p}_i = 0$, it can be seen that the formulae from Equations (S35)-(S36), which describe the steady-state protein mass fraction, still hold for both native and heterologous genes. However, they now include a heterologous protein component:

$$\bar{p}_i = \frac{M}{n_i} \cdot \frac{\bar{m}_i/\bar{k}_i}{\frac{1}{1-\bar{\phi}_q} \sum_{j \in \{a,r\} \cup X} \bar{m}_j/\bar{k}_j} \tag{S62}$$

$$\bar{\phi}_i = \frac{\bar{m}_i/\bar{k}_i}{\frac{1}{1-\bar{\phi}_q} \sum_{j \in \{a,r\} \cup X} \bar{m}_j/\bar{k}_j} \tag{S63}$$

We simulated (Model VI) for a generic heterologous gene, displaying the results in Figure 3 of the main text alongside the analytical predictions for gene expression, which we obtained using the relations derived in the the next three sections of this text. The values of parameters characterizing the gene used are given in Table S4. The same parameters describe the "gene of interest" *poi*, for which we show how to maximise its expression by a population of cells in the Supplementary Information's Section 3.4.

Table S4: Parameters of the generic synthetic genes, the results of simulating which are shown in Figure S2 and the main text's Figure 3.

| Parameter | Description | Value | Units |
|---|---|---|---|
| $c_x = c_{poi}$ | Gene DNA copy number | Varied | $nM$ |
| $\alpha_x = \alpha_{poi}$ | Promoter strength | 1,000 | None |
| $\beta_x = \beta_{poi}$ | mRNA degradation rate | 6 | $h^{-1}$ |
| $k_x^+ = k_{poi}^+$ | mRNA-ribosome binding rate | 60 | $\frac{1}{nM \cdot h}$ |
| $k_x^- = k_{poi}^-$ | mRNA-ribosome dissociation rate | 60 | $nM^{-1}$ |
| $n_x = n_{poi}$ | Number of amino acids in protein $p_i$ | 300 | $aa/nM$ |

#### 3.2 Estimating Steady-State Protein Mass Fractions with Heterologous Expression

The numerical simulations enabled by (Model VI) can be useful in validating gene circuit designs to ensure desired responses despite the influence of resource competition and shifts in the growth rate. Nonetheless, the design of novel resource-aware controllers can also greatly benefit from analytical insights into the system's behavior. Although (Model VI) itself is not analytically tractable, some simplifying assumptions can provide an approximation of the intracellular variables' steady-state values. As demonstrated by simulation results displayed in Figure S2, if the nutrient quality is high, the steady state values of the translation elongation rate $\overline{\epsilon}$ and the ppGpp-dependent ribosomal gene transcription regulation function $\overline{F}_r$ change very little over a wide range of heterologous gene DNA concentrations (and therefore heterologous mRNA production rates). Indeed, Figure S2 shows that even for the highest concentration considered, when the heterologous mRNA production rate exceeds the highest possible combined rate of transcription of all

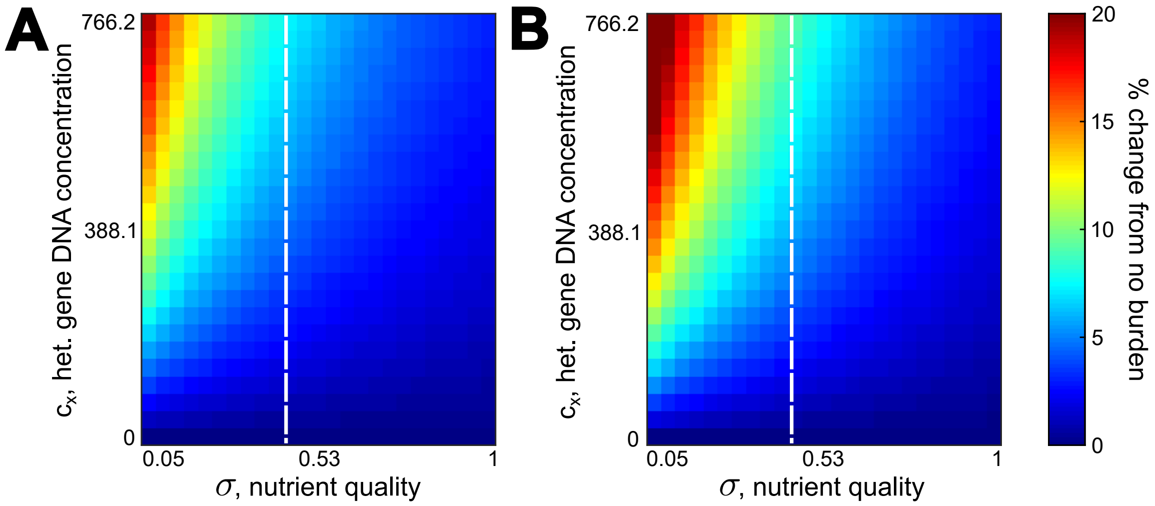

Figure S2: Changes in the cell's steady-state **(A)** translation elongation rate $\overline{\epsilon}$ and **(B)** ribosome transcription regulation function $\overline{F}_r$ caused by expressing a heterologous gene for various heterologous DNA concentrations and culture medium's nutrient qualities. The heterologous gene's parameters are given in Table S4. The white line marks the $\sigma = 0.39$ threshold, right of which $\overline{\epsilon}$ and $\overline{F}_r$ change by no more than 10% even when heterologous mRNA production rate exceeds the maximum possible total rate of transcription of all non-housekeeping native genes.

ribosomal and metabolic genes, neither of these values changes by more than 10% if $\sigma \geq 0.39$. Realistically, heterologous genes are likely to be expressed at a lower rate than in such an extreme case. Hence, the burden-induced changes in $\overline{\epsilon}$ and $\overline{F}_r$ will be smaller than 10% even for culture media of poorer quality.

Therefore, we can assume these two quantities to be independent of the heterologous gene expression burden and the parameters describing heterologous gene expression. Using the index $^{NB}$ to denote the "no burden" steady state values of variables in absence of heterologous gene expression, this assumption can be written as

$$\epsilon \approx \overline{\epsilon}^{NB}, \quad F_r \approx \overline{F}_r^{NB} \quad \forall \{a_{x_l}\}, \{b_{x_l}\}, \{c_{x_l}\}, \{F_{x_l}\} \tag{S64}$$

where $\overline{\epsilon}^{NB}$ and $\overline{F}_r^{NB}$ are easily determined by numerically retrieving the steady state of the host cell expressing no synthetic genes (i.e., simulating (Model III) for a given value of $\sigma$). A corollary to that assumption is that the mRNA-ribosome affinities are also burden-independent, as we show in Equation (S65).

$$k_i = k_i(\epsilon) = \frac{k_i^- + \epsilon/n_i}{k_i^+} \approx \frac{k_i^- + \overline{\epsilon}^{NB}/n_i}{k_i^+} = \overline{k}_i^{NB} \tag{S65}$$

In this case, what is the steady-state mRNA concentration for a given gene, be it native or heterologous? From Equations (S55) and (S60), it follows that

$$\overline{m}_i = \frac{\overline{F}_i c_i \alpha_i \lambda(\overline{\epsilon}^{NB}, \overline{B})}{\beta_i + \lambda(\overline{\epsilon}^{NB}, \overline{B})} \quad \forall i \in \{a, r\} \cup X \tag{S66}$$

where $\overline{F}_i$ is the steady-state value of gene $i$'s regulatory function $F_i$. Provided that the mRNA decay rate is roughly the same for all synthetic genes and native gene classes – that is, $\beta_i \approx \beta_j \ \ \forall i, j \in \{a, r\} \cup X$ – the expression from Equation (S63) for the steady-state protein mass fraction can be rewritten to only include the gene transcription parameters and mRNA-ribosome affinities:

$$\overline{\phi}_i = \frac{\overline{m}_i/\overline{k}_i^{NB}}{\frac{1}{1-\overline{\phi}_q}\sum_{j\in\{a,r\}\cup X}\overline{m}_j/\overline{k}_j^{NB}} = \frac{\frac{\overline{F}_i c_i \alpha_i \lambda(\overline{\epsilon}^{NB}, \overline{B})}{(\beta + \lambda(\overline{\epsilon}^{NB}, \overline{B}))\cdot \overline{k}_i^{NB}}}{\frac{1}{1-\overline{\phi}_q}\sum\limits_{j\in\{a,r\}\cup X}\frac{\overline{F}_j c_j \alpha_j \lambda(\overline{\epsilon}^{NB}, \overline{B})}{(\beta + \lambda(\overline{\epsilon}^{NB}, \overline{B}))\cdot \overline{k}_j^{NB}}} \Leftrightarrow$$

$$\Leftrightarrow \overline{\phi}_i = (1-\overline{\phi}_q)\cdot \frac{\overline{F}_i c_i \alpha_i/\overline{k}_i^{NB}}{\sum\limits_{x_l\in X}\overline{F}_{x_l} c_{x_l}\alpha_{x_l}/\overline{k}_{x_l}^{NB} + \sum\limits_{j\in\{a,r\}}\overline{F}_j c_j \alpha_j/\overline{k}_j^{NB}} \tag{S67}$$

Importantly, under the assumption of constant translation elongation rate and unchanging ppGpp signal regulating ribosomal gene transcription, Equation (S67)'s terms for both ribosomal and metabolic genes, which together constitute the set $\{a, r\}$, are constant and independent of the identity and parameters of the heterologous genes being expressed. Let us then define $\xi$, the "translational burden" or "translational demand" of heterologous gene expression – that is, the extent of competition for ribosomes collectively imposed by all synthetic genes. This can be found by calculating the product of the copy number, promoter strength and mRNA-ribosome dissociation constant for every heterologous gene and adding these quantities together, as shown in Equation (S68).

$$\xi = \sum_{x_l \in X} \frac{\overline{F}_{x_l} c_{x_l} \alpha_{x_l}}{\overline{k}^{NB}_{x_l}} \tag{S68}$$

Consequently, the total mass fraction of all heterologous genes $\overline{\phi}_X$ follows a relationship akin to the Hill activation function:

$$\overline{\phi}_X(\xi) = \sum_{x_l \in X} \overline{\phi}_{x_l} = \frac{\xi}{\xi + (\sum_{j \in \{a,r\}} \overline{F}^{NB}_j c_j \alpha_j / \overline{k}^{NB}_j)} \tag{S69}$$

The mass fraction of an individual gene is therefore given by:

$$\overline{\phi}_{x_l}(\xi) = \overline{\phi}_X(\xi) \cdot \frac{\overline{F}_{x_l} c_{x_l} \alpha_{x_l} / \overline{k}^{NB}_{x_l}}{\sum_{j \in X} \overline{F}_j c_j \alpha_j / \overline{k}^{NB}_j} = \overline{\phi}_X(\xi) \cdot \frac{\overline{m}_{x_l} / \overline{k}_{x_l}}{\sum_{j \in X} \overline{m}_j / \overline{k}_j} \qquad \forall x_l \in X \tag{S70}$$

As for the native genes $a$ and $r$, substitution into Equation (S67) shows that their mass fractions, conversely, obey a Hill repressor function-like law:

$$\overline{\phi}_i(\xi) = (1 - \overline{\phi}_q) \cdot \frac{\overline{F}^{NB}_i c_i \alpha_i / \overline{k}^{NB}_i}{\xi + \sum_{j \in \{a,r\}} \overline{F}_j c_j \alpha_j / \overline{k}^{NB}_j} \qquad \forall i \in \{a, r\} \tag{S71}$$

#### 3.3 Estimating Steady-State Growth Rate with Heterologous Gene Expression

From the previous section, we have an intuition how the steady-state protein mass fractions are affected by the translational burden of expressing heterologous genes. We now use this knowledge to find how burden influences cell growth. Let us recall the definition of growth rate $\lambda$ from Table S2. In steady state, it yields:

$$\overline{\lambda} = \lambda(\overline{\epsilon}^{NB}, \overline{B}) = \frac{\overline{\epsilon}^{NB} \cdot \overline{B}}{M}$$

Let us assume that the share of free ribosomes is negligible – indeed, even without heterologous mRNA transcription, at high nutrient qualities $D$ is very high (on the order of $10^4$), so the share of ribosomes not engaged in translation, $\frac{1}{1+D}$, is insignificant. Then, we have $B \approx R$, which transforms the expression for the steady-state growth rate into

$$\overline{\lambda} \approx \frac{\overline{\epsilon}^{NB} \cdot \overline{R}}{M} \Leftrightarrow$$

$$\Leftrightarrow \overline{\lambda}(\xi) = \frac{\overline{\epsilon}^{NB}}{n_r} \cdot \overline{\phi}_r(\xi) = \frac{\overline{\epsilon}^{NB}(1-\overline{\phi}_q)}{M} \cdot \frac{\overline{F}_r^{NB} c_r \alpha_r / \overline{k}_r^{NB}}{\xi + \sum\limits_{j\in\{a,r\}} \overline{F}_j^{NB} c_j \alpha_j / \overline{k}_j^{NB}} \tag{S72}$$

This is similar in form to the Hill relationship between the heterologous gene expression burden and the steady-state growth rate observed by McBride et al. [1]. Meanwhile, the ratio between the growth rates with and without heterologous gene expression burden can be found as shown in Equation (S73). Meanwhile, this matches the linear-like relationship between the growth rate and heterologous protein mass fraction observed by Scott et al. [8].

$$\frac{\overline{\lambda}(\xi)}{\overline{\lambda}^{NB}} \approx \left( \frac{\overline{\epsilon}^{NB}(1-\overline{\phi}_q)}{M} \cdot \frac{\overline{F}_r^{NB} c_r \alpha_r / \overline{k}_r^{NB}}{\xi + \sum\limits_{j\in\{a,r\}} \overline{F}_j^{NB} c_j \alpha_j / \overline{k}_j^{NB}} \right) \div \left( \frac{\overline{\epsilon}^{NB}(1-\overline{\phi}_q)}{M} \cdot \frac{\overline{F}_r^{NB} c_r \alpha_r / \overline{k}_r^{NB}}{\sum\limits_{j\in\{a,r\}} \overline{F}_j^{NB} c_j \alpha_j / \overline{k}_j^{NB}} \right) \Leftrightarrow$$

$$\Leftrightarrow \frac{\overline{\lambda}(\xi)}{\overline{\lambda}^{NB}} \approx \frac{\sum\limits_{j\in\{a,r\}} \overline{F}_j^{NB} c_j \alpha_j / \overline{k}_j^{NB}}{\xi + \sum\limits_{j\in\{a,r\}} \overline{F}_j^{NB} c_j \alpha_j / \overline{k}_j^{NB}} = 1 - \frac{\xi}{\xi + \sum\limits_{j\in\{a,r\}} \overline{F}_j^{NB} c_j \alpha_j / \overline{k}_j^{NB}} \Leftrightarrow$$

$$\Leftrightarrow \frac{\overline{\lambda}(\xi)}{\overline{\lambda}^{NB}} \approx 1 - \frac{\overline{\phi}_X(\xi)}{1-\overline{\phi}_q} \tag{S73}$$

#### 3.4 Maximizing Heterologous Protein Production

A prominent application of synthetic biology is bioproduction of valuable proteins [14]. In order to ensure the process's cost-efficiency, it is thus desirable to maximise the yield of the protein produced by a population of cells in a bioreactor. A key hindrance to achieving this, however, is the need to balance high protein synthesis rates with small gene expression burden. Indeed, while a single cell can be made to produce a lot of protein, severe impairment of growth by high heterologous gene expression would mean that the cell population expands very slowly, so the overall rate of protein production would in fact be low [15].

The relations derived in Section 3.2 and 3.3 can be leveraged to address this issue. Let us define a simple model of a homogenous population of *E. coli* cells, dying at a constant rate $\delta$ [2], where $N$ denotes the overall number of cells, whereas $\lambda$ is the cell growth rate as predicted by our model (Equation (S74)).

$$\dot{N} = \lambda N - \delta N \tag{S74}$$

If the bacteria express a single gene *poi* encoding the intracellular protein of interest and are in steady state, how would we approach maximizing the protein yield? Protein production per cell is commonly characterized as the product of the amount of protein in one cell and the rate at which the population expands [16]. Under our model, this is equal to:

$$\mu = M \cdot \overline{\phi}_{poi} \cdot (\overline{\lambda} - \delta) \tag{S75}$$

Since the steady-state growth rate can be estimated with the help of Equation (S73), we can make a substitution to obtain:

$$\mu \approx M \cdot \overline{\phi}_{poi} \cdot \left( \overline{\lambda}^{NB} \left( 1 - \frac{\overline{\phi}_{poi}}{1 - \overline{\phi}_q} \right) - \delta \right) \Leftrightarrow$$

$$\Leftrightarrow \mu \approx - \frac{\overline{\lambda}^{NB}}{1 - \overline{\phi}_q} M \overline{\phi}^2_{poi} + M(\overline{\lambda}^{NB} - \delta)\overline{\phi}_{poi} \tag{S76}$$

This is a quadratic expression, so the value of $\overline{\phi}_{poi}$ that maximizes it can be easily calculated as:

$$\overline{\phi}^{max}_{poi} = \frac{1}{2} \left(1 - \overline{\phi}_q\right) \left(1 - \frac{\delta}{\overline{\lambda}^{NB}}\right) \tag{S77}$$

Knowing this value, we can use Equation (S69) to find the translational burden $\xi_{max}$ corresponding to the optimal heterologous protein mass fraction. This allows to ensure maximum protein production by imposing a condition on the synthetic gene's mRNA-ribosome dissociation constant, gene copy number and promoter strength, which we provide below.

$$\begin{aligned} \overline{\phi}_{poi}(\xi_{max}) = \overline{\phi}_{poi}^{max} \Leftrightarrow \\ \Leftrightarrow \frac{c_{poi}\alpha_{poi}}{\overline{k}_{poi}^{NB}} = \xi_{max} = \frac{1-\delta/\overline{\lambda}^{NB}}{1+\delta/\overline{\lambda}^{NB}} \cdot \sum_{j\in\{a,r\}} \frac{\overline{F}_j c_j \alpha_j}{\overline{k}_j^{NB}} \end{aligned} \tag{S78}$$

### 4 Biocircuit modeling

In this section, we define the ODEs for the synthetic gene circuits considered in the article. We also provide derivations of the analytical relations that can guide biocircuit design and show the results of simulations that were conducted in addition to those described in the main text.

#### 4.1 Two Bistable Switches Exhibiting "Winner-Takes-All" Behavior

##### 4.1.1 Description

To demonstrate that our model captures known resource competition phenomena, we considered the case of two synthetic self-activating genes being present in the same cell. Here, "self-activation" means that the protein encoded by a gene can bind a corresponding inducer molecule to form a complex that acts as a transcription activation (TA) factor for the same gene. On its own, such a gene can act as a bistable switch due to having two different stable steady states, one with high gene expression and the other with low expression. Meanwhile, two switches in the same cell may exhibit "winner-takes-all" behavior – that is, if one gene reaches its high-expression equilibrium faster than the other, the increased resource competition from it can prevent the second gene from becoming highly expressed [17].

For simplicity, we characterized the two synthetic self-activating genes $s_1$ and $s_2$ by the same parameter values, displayed in Table S5. The ODEs describing their expression mirrored the generic Equations (S60) and (S61) for heteroglous mRNA and protein concentrations, hence:

$$\dot{m_{s_1}} = F_{s_1}(p_{s_1}) \cdot c_{s_1}\alpha_{s_1} - (\beta_{s_1} + \lambda(\epsilon, B))m_{s_1} \tag{S79}$$

$$\dot{p_{s_1}} = \frac{\epsilon(t^c)}{n_{s_1}} \cdot \frac{m_{s_1}/k_{s_1}}{1 + \frac{1}{1-\bar{\phi}_q}\sum_{j\in\{a,r\}\cup X} m_j/k_j} R - \lambda(\epsilon, B) \cdot p_{s_1} \tag{S80}$$

$$\dot{m_{s_2}} = F_{s_2}(p_{s_2}) \cdot c_{s_2}\alpha_{s_2} - (\beta_{s_2} + \lambda(\epsilon, B))m_{s_2} \tag{S81}$$

$$\dot{p_{s_2}} = \frac{\epsilon(t^c)}{n_{s_2}} \cdot \frac{m_{s_2}/k_{s_2}}{1 + \frac{1}{1-\bar{\phi}_q}\sum_{j\in\{a,r\}\cup X} m_j/k_j} R - \lambda(\epsilon, B) \cdot p_{s_2} \tag{S82}$$

where the transcription regulation function for gene $s_i$ is defined as:

$$F_{s_i}(f_i, p_{s_i}) = \frac{F_{s_i,0} \cdot K_{dna_i}^{\eta_i} + \left( \frac{f_i}{K_{ind_i} + f_i} \cdot p_{s_i} \right)^{\eta_i}}{K_{dna_i}^{\eta_i} + \left( \frac{f_i}{K_{ind_i} + f_i} \cdot p_{s_i} \right)^{\eta_i}} \tag{S83}$$

Here, $f_i$ is the inducer concentration. To simulate the activation of a switch at time $t_{induction}$, we made it follow a step function of time displayed below. In order to investigate different possible behaviors of the system, different combinations of the two genes' added inducer concentrations, $f_1^{added}$ and $f_2^{added}$, were considered as outlined in the main text.

$$f_i(t) = \begin{cases} 0, \text{ if } t < t_{induction} \\ f_i^{added}, \text{ otherwise} \end{cases} \tag{S84}$$

Table S5: Parameters of the synthetic self-activating genes.

| Parameter | Description | Value | Units |
|---|---|---|---|
| $c_{s_i}$ | Gene copy number | 100 | $nM$ |
| $\alpha_{s_i}$ | Promoter strength | 3,000 | None |
| $\beta_{s_i}$ | mRNA degradation rate | 6 | $h^{-1}$ |
| $k_{s_i}^+$ | mRNA-ribosome binding rate | 60 | $\frac{1}{nM \cdot h}$ |
| $k_{s_i}^-$ | mRNA-ribosome dissociation rate | 60 | $nM^{-1}$ |
| $n_{s_i}$ | Number of amino acids in protein $p_i$ | 300 | $aa/nM$ |
| | **Gene transcription regulation function** | | |
| $F_{s_i,0}$ | Baseline function value in absence of inducer | 0.1 | None |
| $K_{ind_i}$ | Half-saturation constant for protein-inducer binding | 500 | $\mu M$ |
| $K_{dna_i}$ | Half-saturation constant for the TA factor-promoter DNA binding | 50,000 | $nM$ |
| $\eta_i$ | Cooperativity coefficient for the TA factor-promoter DNA binding | 2 | $nM$ |

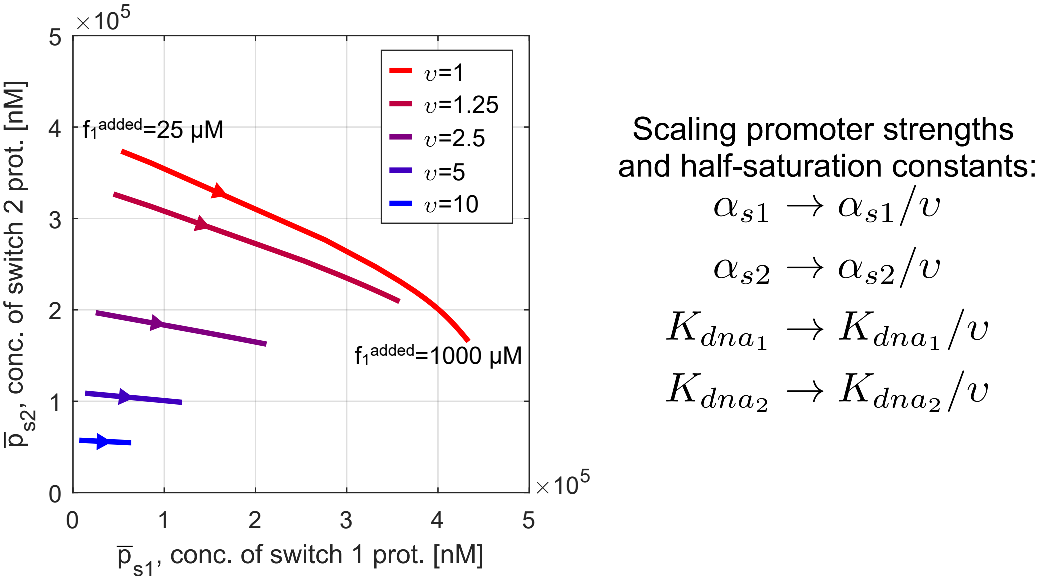

Figure S3: Phase plane diagram for the system of two self-activating bistable switch genes with the promoter strengths and DNA-transcription factor binding half-saturation constants scaled by $^1/_\upsilon$ for different values of $\upsilon$. Bold lines show the system's steady states in presence of 125 $nM$ of inducer 2 and concentrations of inducer 1 ranging between 25 $nM$ and 1000 $nM$. The arrows on the lines point in the direction of increasing inducer 1's concentration, $f_1$.

##### 4.1.2 Additional Simulations

For a system of two self-activating bistable switch genes in the same host cell, additional simulations were performed besides those discussed in the main text's Results section and displayed in the main text's Figure 3F. The details of these numerical experiments are outlined in this section.

In the main text, we discussed the "winner-takes-all" phenomenon exhibited by the described system, where the switch that reaches a high-expression equilibrium first (the "winner") prevents the other switch from becoming activated due to resource couplings. In Figure 3F, we demonstrated this by varying the concentration of the first self-activating gene's inducer, $f_1$. Since the gene with a higher inducer concentration becomes activated first, setting $f_1$ lower than that the second gene's inducer concentration $f_2$ led to the second switch "winning". Conversely, for $f_1 > f_2$ the first switch gene prevailed while the expression of the second remained low.

However, in order for this to occur, resource competition exerted by the switches must be great enough to significantly affect gene expression. To investigate the relationship between "winner-takes-all" behavior and the burden caused by the switch genes, we repeated the simulation outlined above but scaled the genes' promoter strengths by a factor of $^1/_\upsilon$, since a weaker promoter means that fewer mRNA molecules are transcribed, so fewer transcripts compete for ribosomes. As decreasing transcription also reduces the overall amount of protein produced, the half-saturation constants for DNA-transcription factor binding were scaled, too – otherwise, $p_{s_i}$ could be too low compared to $K_{dna_i}$ for the switch to exhibit bistability. All other parameters remained as given in Table S5.

Figure S3 shows that as $\upsilon$ increases, the effect of higher inducer 1 concentrations on the second gene's expression becomes less significant. Accordingly, when $\upsilon = 10$, the line that spans the system's equilibria for $25\ \mu M < f_1 < 1000\ \mu M$ becomes almost horizontal, indicating that the steady-state value of $p_2$ is minimally affected. Numerical simulations conducted with the help of our model can therefore not only capture the well-known resource competition phenomenon "winner-takes-all" behavior, but also allow to determine if it will be exhibited for a given combination of design parameters. However, it is important to note that *in vivo* a switch circuit is affected not only by resource couplings but also by other factors, such as the stochasticity of gene expression, which can also significantly alter its behavior [18].

#### 4.2 Proportional-Integral Controller for Resource Coupling Mitigation

##### 4.2.1 Description

Here, we consider the antithetic integral feedback controller that maintains a constant level of competition for ribosomes in the cell and thus a constant growth rate. This circuit's working principle is outlined in the main text's Results section.

Briefly, it consists of four genes: the "sensor" *sens*, the "actuator" *act*, the "annihilator" *anti* and the "amplifier" *amp*. The actuator is a constitutive gene, whose mRNA $m_{act}$ can be translated to produce the regulatory protein $p_{act}$ that activates the expression of the amplifier gene's mRNA $m_{amp}$. As the amplifier gene's maximum transcription rate is chosen to be high, $m_{amp}$ significantly contributes to competition for the cell's translational resources by binding ribosomes in order to be translated into a non-toxic protein $p_{amp}$. Meanwhile, the sensor gene is constitutive and encodes a repressor protein $p_{sens}$, which regulates the transcription of the annihilator RNA $m_{anti}$. Since $m_{anti}$ is an antisense RNA strand binding and inactivating $m_{act}$, it affects the production of $m_{amp}$. Therefore, the cell's resource competition landscape is manipulated in response to resource coupling-induced changes in the expression of the constitutively expressed protein $p_{sens}$. Consequently, if we assume that the binding of all regulatory proteins occurs non-cooperatively, the transcription regulation functions are as follows:

$$F_{sens} \equiv F_{act} \equiv 1, \quad F_{anti} = \frac{K_{sens}}{K_{sens} + p_{sens}}, \quad F_{amp} = \frac{p_{act}}{K_{act} + p_{act}} \tag{S85}$$

The ODEs describing the system are given in Equations (S86)-(S92). As for the model's parameters, different genes' copy numbers and maximum transcription rates $c_i$ and $\alpha_i$ can be multiplied together as shown in Table S6 to yield quantities that are more informative of the controller's behavior. We thus use the parameters $\zeta, u, \kappa$ and $\chi$ in model equations and in Table S7 displaying the values of the model parameters.

$$\dot{m}_{sens} = \zeta\lambda(\epsilon, B) - (\beta_{sens} + \lambda(\epsilon, B))m_{sens} \tag{S86}$$

$$\dot{m}_{anti} = F_{anti}\kappa\lambda(\epsilon, B) - (\beta_{anti} + \lambda(\epsilon, B))m_{anti} - \theta m_{act}m_{anti} \tag{S87}$$

$$\dot{m}_{act} = u\kappa\lambda(\epsilon, B) - (\beta_{act} + \lambda(\epsilon, B))m_{act} - \theta m_{act}m_{anti} \tag{S88}$$

$$\dot{m}_{amp} = F_{amp}\chi\lambda(\epsilon, B) - (\beta_{amp} + \lambda(\epsilon, B))m_{amp} \tag{S89}$$

$$\dot{p}_{sens} = \frac{\epsilon(t^c)}{n_{sens}} \cdot \frac{m_{sens}/k_{sens}}{1 + \frac{1}{1-\bar{\phi}_q}\sum_{j\in\{a,r\}\cup X} m_j/k_j} R - \lambda(\epsilon, B) \cdot p_{sens} \tag{S90}$$

$$\dot{p}_{act} = \frac{\epsilon(t^c)}{n_{act}} \cdot \frac{m_{act}/k_{act}}{1 + \frac{1}{1-\bar{\phi}_q}\sum_{j\in\{a,r\}\cup X} m_j/k_j} R - \lambda(\epsilon, B) \cdot p_{act} \tag{S91}$$

$$\dot{p}_{amp} = \frac{\epsilon(t^c)}{n_{amp}} \cdot \frac{m_{amp}/k_{amp}}{1 + \frac{1}{1-\bar{\phi}_q}\sum_{j\in\{a,r\}\cup X} m_j/k_j} R - \lambda(\epsilon, B) \cdot p_{amp} \tag{S92}$$

We are also interested in observing how the controller mitigates external disturbances in the form of additional competition form other heterologous genes' mRNAs. Here, we investigate the simplest-possible case of one additional heterologous gene *dist*, whose transcription is controlled by a function $F_{dist}$ that rises linearly from 0 (no disturbance) to 1 (full expression of disturbing mRNA) over a brief period of 0.1 $h$ and then stays at $F_{dist} = 1$. This setup models an abrupt increase in the number of mRNAs competing for ribosomes, which the AIF controller is expected to counter by decreasing the concentration of another competing species $m_{act}$. Since the disturbing gene's transcript and protein are not involved in any additional reactions besides trancription, translation, dilution and mRNA degradation, ODEs describing the dynamics of the corresponding mRNA concentration $m_{dist}$ and protein concentration $p_{dist}$ are trivial and can be readily obtained from (Model VI)'s Equations (S60)-(S61) by replacing the generic $x_l$ index with *dist*.

When in the main text's Figure 5H-I we consider the case of two synthetic genes, *dist* and $x$, being expressed besides our controller, the expression of the constitutive "output" gene $x$ is likewise straightforwardly described by replacing the generic index $x_l$ with $x$ in (Model VI)'s Equations (S60)-(S61) and postulating $F_x \equiv 1$.

##### 4.2.2 Circuit Analysis

The error reacted upon by the antithetic integral feedback is the difference between $u$ and $F_{act}$, which is determined by the sensor protein concentration $p_{sens}$. However, knowing the circuit parameter values, we can use the derivations from Section 3 to calculate from circuit parameters the values of the resource competition denominator $\overline{D}$ (i.e., the variable we actually desire to keep constant) and the growth rate $\overline{\lambda}$ enforced by the controller. Furthermore, it is possible to find the controller's operation range boundary – that is, the maximum gene expression burden that it can mitigate.

The dilution and degradation of $m_{act}$ and $m_{anti}$ lead to the phenomenon of "leakiness", which prevents the antithetic integral feedback controller from reaching perfect adaptation. In the idealized case, however, it can be considered negligible [19]. Making this assumption in order to obtain easily calculable analytical estimates for $\overline{D}$ and $\overline{\lambda}$, we obtain the following steady-state relations:

$$\begin{cases} \dot{m}_{act} \approx \overline{\lambda}\kappa u - \theta \overline{m}_{act}\overline{m}_{anti} = 0 \\ \dot{m}_{anti} \approx \overline{\lambda}\kappa F_{anti} - \theta \overline{m}_{act}\overline{m}_{anti} = 0 \end{cases} \Rightarrow \kappa u = \kappa F_{anti} \Leftrightarrow$$

$$\Leftrightarrow \overline{\phi}_{sens} = \frac{n_{sens}}{M} \cdot K_{sens} \cdot \frac{1-u}{u} \tag{S93}$$

According to Equation (S63), it can thus be observed that

$$\frac{n_{sens}}{M} \cdot K_{sens} \cdot \frac{1-u}{u} = \frac{\overline{m}_{sens}/\overline{k}^{NB}_{sens}}{\overline{D}-1} \Leftrightarrow$$

$$\Leftrightarrow \overline{D} - 1 = \frac{\overline{m}_{sens}}{\overline{k}^{NB}_{sens}} \cdot \left( \frac{n_{sens}}{M} \cdot K_{sens} \cdot \frac{1-u}{u} \right)^{-1} \tag{S94}$$

Then, by assuming that the share of idle ribosomes is negligible (likewise to Section 3.3), we obtain:

$$\overline{\lambda} = \frac{\overline{\epsilon}^{NB}}{M}\overline{B} \approx \frac{\overline{\epsilon}^{NB}}{M}\overline{R} \Leftrightarrow \overline{\lambda} \approx \frac{\overline{\epsilon}^{NB}}{M} \cdot \frac{M}{n_r} \cdot \frac{\overline{m}_r/\overline{k}_r^{NB}}{\overline{D}-1} \Leftrightarrow$$
$$\Leftrightarrow \overline{\lambda} \approx \frac{\overline{\epsilon}^{NB}}{n_r} \cdot \frac{\overline{m}_r}{\overline{k}_r^{NB}} \cdot \left(\frac{\overline{m}_{sens}}{\overline{k}_{sens}^{NB}}\right)^{-1} \cdot \frac{n_{sens}}{M} \cdot K_{sens} \cdot \frac{1-u}{u} \tag{S95}$$

Now, let us use substitute the formula for steady-state mRNA concentrations from Equation (S66) for the sensor and ribosomal gene transcripts. Assuming the two mRNAs to have similar degradation rates, we make the following simplification:

$$\frac{\overline{m}_r}{\overline{m}_{sens}} = \frac{\overline{F}_r^{NB} c_r \alpha_r \overline{\lambda}}{\overline{\lambda} + \beta_r} \cdot \left(\frac{\zeta\overline{\lambda}}{\overline{\lambda} + \beta_{sens}}\right)^{-1} = \frac{\overline{F}_r^{NB} c_r \alpha_r}{\zeta} \tag{S96}$$

Substituting Equation (S96) into Equation (S95), we finally obtain a formula for the steady-state growth rate of the cell:

$$\overline{\lambda} \approx \frac{\overline{\epsilon}^{NB}}{M} \cdot \overline{F}_r^{NB} c_r \alpha_r \cdot \frac{n_{sens}\overline{k}_{sens}^{NB}}{n_r \overline{k}_r^{NB}} \cdot \frac{K_{sens}}{\zeta} \cdot \frac{1-u}{u} \tag{S97}$$

Knowing the growth rate, from Equation (S86) we can find the steady-state value of the sensor gene mRNA as

$$\overline{m}_{sens} = \frac{\overline{\lambda}\zeta}{\overline{\lambda} + \beta_{sens}} \tag{S98}$$

which in turn can be plugged into Equation (S94) to yield an estimate for the value of $\overline{D}$ — that is, the extent of resource competition maintained by the controller:

$$\overline{D} = 1 + \frac{\overline{\lambda}\zeta}{(\overline{\lambda} + \beta_{sens})\overline{k}_{sens}^{NB}} \cdot \left(\frac{n_{sens}}{M} \cdot K_{sens} \cdot \frac{1-u}{u}\right)^{-1} \tag{S99}$$

Moreover, with the growth rate known, we can find the total steady-state level of the actuator and amplifier mRNAs. This is useful because the controller combats the effects of the appearance of extra competing heterologous mRNAs by decreasing $m_{amp}$. The abundance of the actuator mRNA $m_{act}$ is likewise decreased in this case, albeit this has more relevance for regulating the amplifier gene's expression via $p_{act}$ levels rather than for its direct contribution to resource competition, which is small relatively to that of $m_{amp}$. The maximum possible total concentrations of $m_{amp}$ and $m_{act}$ are thus achieved in the case of no heterologous gene expression. Meanwhile, since the actuator mRNA level cannot be decreased below zero, the value of

$$max\left(\frac{m_{act}}{k_{act}} + \frac{m_{amp}}{k_{amp}}\right)$$

defines the controller's operation range. By recalling Equations (S93), (S63) and (S66), we can find this sum for the case of no heterologous genes being expressed besides those comprizing the controller. Yielded by the formula in Equation (S100), it gives rise to the definition of our controller's operating range in the main text.

$$\frac{n_{sens}}{M} \cdot K_{sens} \cdot \frac{1-u}{u} = (1-\overline{\phi}_q) \cdot \frac{\overline{m}_{sens}/\overline{k}^{NB}_{sens}}{\overline{m}_{act}/\overline{k}^{NB}_{act} + \overline{m}_{amp}/\overline{k}^{NB}_{amp} + \sum_{j\in\{a,r,sens\}} \overline{m}_j/\overline{k}^{NB}_j} \Leftrightarrow$$

$$\Leftrightarrow \frac{\overline{m}_{act}}{\overline{k}^{NB}_{act}} + \frac{\overline{m}_{act}}{\overline{k}^{NB}_{act}} = \frac{(1-\overline{\phi}_q)M\zeta\overline{\lambda}}{K_{sens}n_{sens}\overline{k}^{NB}_{sens}(\overline{\lambda}+\beta_{sens})} \cdot \frac{1-u}{u} - \sum_{j\in\{a,r,sens\}} \frac{\overline{F}^{NB}_j c_j \alpha_j \overline{\lambda}}{\overline{k}^{NB}_j(\overline{\lambda}+\beta_j)} \quad \text{(S100)}$$

Table S6: Calculation of informative model parameters from gene copy numbers and promoter strengths.

| Variable | Description | Formula | Units |
|---|---|---|---|
| $\zeta$ | Total transcription rate of sensor mRNA | $c_{sens}\alpha_{sens}$ | $nM^*$ |
| $\kappa$ | Maximum total transcription rate of annihilator RNA | $c_{anti}\alpha_{anti}$ | $nM^*$ |
| $u$ | Value of $F_{anti}$ enforced by the controller | $\frac{c_{act}\alpha_{act}}{c_{anti}\alpha_{anti}}$ | None |
| $\chi$ | Maximum total transcription rate of amplifier RNA | $c_{amp}\alpha_{amp}$ | $nM^*$ |

$^*$ Transcription rates given in the units of $nM$ because in Equations (S86)-(S89) the mRNA production rate is calculated by multiplying these parameters by the cell growth rate $\lambda$ (units of $h^{-1}$).

Table S7: Parameters for simulating the synthetic gene circuit described by Equations (S88)-(S90).

| Parameter | Description | Value | Units |
|---|---|---|---|
| $\kappa$ | Max. total transc. rate of annihilator RNA | $2 \cdot 10^5$ | $nM$ |
| $u$ | Value of $F_{anti}$ enforced by the controller | 0.5 | None |
| $\zeta$ | Total transc. rate of sensor mRNA | $1.5 \cdot 10^4$ | $nM$ |
| $\theta$ | Actuator-annihilator binding rate | 300 | $\frac{1}{nM \cdot h}$ |
| $K_{sens}$ | Half-saturation constant for $p_{sens}$-annihilator promoter DNA binding | $4,000$ | $nM$ |
| $\eta$ | Hill coefficient for $p_{sens}$-annihilator promoter DNA binding | 1 | None |
| $\beta_i$ | mRNA degradation rate* | 6 | $h^{-1}$ |
| $k_i^+$ | mRNA-ribosome binding rate‡ | 60 | $\frac{1}{nM \cdot h}$ |
| $k_i^-$ | mRNA-ribosome dissociation rate‡ | 60 | $nM^{-1}$ |
| $n_i$ | Number of amino acids in protein $p_i$ ‡ | 300 | $aa/nM$ |
| | **Disturbance gene** | | |
| $c_{dist}$ | Disturbing gene's copy number | 100 | $nM$ |
| $\alpha_{dist}$ | Max. transcription rate for disturbing gene | 300 | $h^{-1}$ |
| | **Output gene** | | |
| $c_x$ | Output gene's copy number | 100 | $nM$ |
| $\alpha_x$ | Max. transcription rate for output gene | 100 | $h^{-1}$ |
| | **Culture medium** | | |
| $\sigma$ | Culture medium's nutrient quality | 0.5 | None |

‡ Identical for all protein-encoding genes *act*, *sens*, *dist* and $x$. Not defined for the annihliator gene *anti*, as it is not translated.

* Identical for all genes.

##### 4.2.3 Additional simulations

Here, we provide the outcomes of additional simulations of our controller's behavior besides those given in the main text's Results section. Likewise to the numerical experiments described in the main text, all circuit parameters were taken from Table S7 unless stated otherwise.

As we point out in the main text's Results and Discussion sections, the proportional feedback component, arising from the dependence of $p_{act}$ synthesis rate on ribosome availability, helps combat the adaptation error (i.e., the difference between the controlled variable's steady-state values before and after perturbation) caused by leakiness. To validate this notion, we considered the hypothetical case of our controller having no proportional feedback. This was achieved by redefining the ODE for the actuator protein as

$$\dot{p}_{act} = \frac{\epsilon(t^c)}{n_{act}} \cdot m_{act} \cdot 0.0660 - \lambda(\epsilon, B) \cdot p_{act} \quad \text{(S101)}$$

instead of the expression from Equation (S91), where 0.0660 is equal to the steady-state value of $\frac{R}{k_{act}D}$ for the case when the controller is the only synthetic gene circuit in the cell. We then repeated the simulation displayed in Figure 5B–F of the main text, where the system was disturbed by the induction of an additional heterologous gene *dist*. The outcome, shown in Figure S4A–E, demonstrated that removing the proportional component slightly increased the adaptation errors for the controlled variable $D$ and the cell growth rate $\lambda$. Namely, for $D$ the error rose to 0.16% from 0.15% observed in presence of proportional feedback. Meanwhile, for $\lambda$ the adaptation error amounted to 1.20% without proportional feedback compared to 1.15% for the full proportional-integral controller.

Moreover, in the main text's Results section we mentioned that increasing the actuator and the annihilator genes' synthesis and mutual annihilation rates – that is, $\kappa$ and $\theta$ – can arbitrarily reduce the error between the steady-state value of the controlled variable and its setpoint [19]. However, this can also lead to instability, although the variable's *average* value will keep converging to the setpoint as the rates increase [20, 21]. As an example of this behavior, in Figure S4F–J we simulate the behavior of a controller whose parameters $\kappa$ and $\theta$ are ten times the values given in Table S7 (while all other parameters' values remain the same). For consistency with Figure S4A–E and the main text's Figure 5, we perturb the system by inducing the expression of an additional heterologous gene *dist* here, too. Notably, while the system exhibits oscillation, the difference between our analytical estimates for $\lambda$ and $D$ and the average values maintained by the controller was smaller for the original design. For $\lambda$, it decreased from 7.69% to only 1.68%, while for $D$ it shrank from 2.0% to 1.82%.

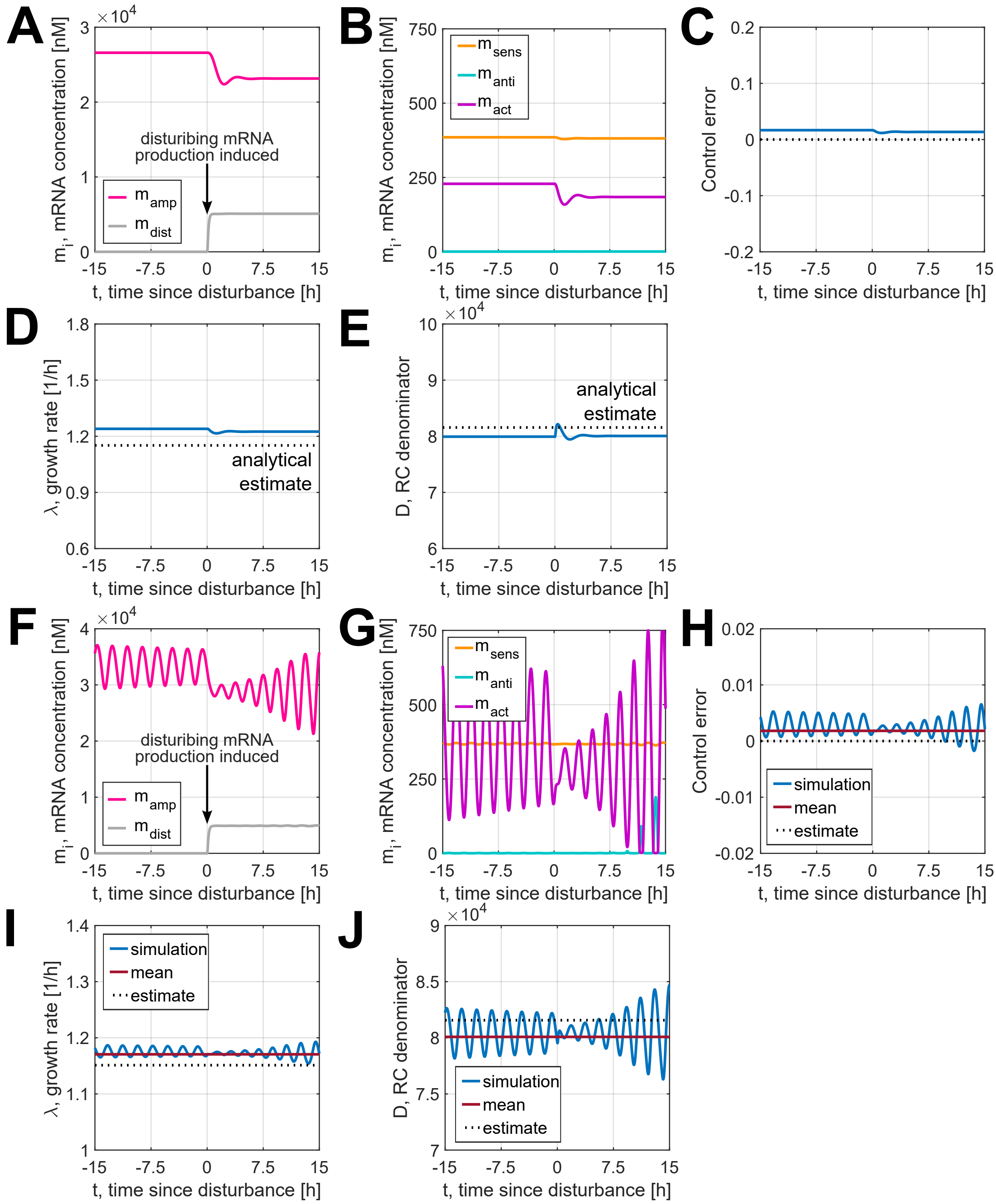

Figure S4: Behavior of modified controllers upon a change in burden due to the induction of an additional synthetic gene *dist*. **(A-E)** The integral-only controller with proportional feedback removed in line with Equation (S101). Figures (A) and (B) display the evolution of mRNA concentrations over time, while figures (C),(D),(E) show the control error sensed by the AIF motif, the cell's growth rate and the resource competition denominator $D$, which is the circuit's controlled variable. **(F-J)** The proportional-integral controller with $\kappa$ and $\theta$ increased tenfold. Figures (F) and (G) display the evolution of mRNA concentrations over time, while figures (H),(I),(J) show the control error sensed by the AIF motif, the cell's growth rate and the resource competition denominator $D$, which is the circuit's controlled variable.

[5] Ron Milo and Rob Phillips. *Cell biology by the numbers*. Garland Science, Abingdon, 2015.

[14] Germán L. Rosano and Eduardo A. Ceccarelli. Recombinant protein expression in escherichia coli: advances and challenges. *Frontiers in Microbiology*, 5, 2014.

[15] Francesca Ceroni, Rhys Algar, Guy-Bart Stan, and Tom Ellis. Quantifying cellular capacity identifies gene expression designs with reduced burden. *Nature Methods*, 12(5):415–418, 2015.

[16] Duncan Ingram and Guy-Bart Stan. When synthetic biology fails: a modular framework for modelling genetic stability in engineered cell populations. *bioRxiv*, 2022.
